# Pervasive transcription-dependent chromatin remodeling influences the replication initiation program

**DOI:** 10.1101/272666

**Authors:** Julien Soudet, Jatinder Kaur, Françoise Stutz

## Abstract

In Eukaryotic organisms, replication initiation follows a temporal program. Among the parameters that regulate this program in *Saccharomyces cerevisiae*, chromatin structure has been at the center of attention without considering the contribution of transcription. Here, we revisit the replication initiation program in the light of pervasive transcription. We find that noncoding RNA transcription termination in the vicinity of replication origins or ARS (Autonomously Replicating Sequences) maximizes replication initiation by restricting transcriptional readthrough into ARS. Consistently, high natural nascent transcription correlates with low ARS efficiency and late replication timing. High readthrough transcription is also linked to chromatin features such as high levels of H3K36me3 and deacetylated nucleosomes. Moreover, forcing ARS readthrough transcription promotes these histone modifications. Finally, replication initiation defects induced by increased transcriptional readthrough are partially rescued in the absence of H3K36 methylation. Altogether, these observations indicate that natural pervasive transcription into ARS influences replication initiation through chromatin remodeling.

## INTRODUCTION

DNA replication is one of the fundamental processes occurring in all living organisms and ensuring accurate duplication of the genome. Eukaryotic replication initiation takes place at several dispersed locations termed replication origins. Origins are defined by a specific chromatin structure consisting of a nucleosome-depleted region (NDR) and the binding of specific replication initiation factors. In *Saccharomyces cerevisiae*, replication origins or ARS (Autonomously Replicating Sequences) are specified by an 11bp T-rich ARS consensus sequence (ACS)^1,2^. ARS also contain more degenerate A-rich B elements proposed to contribute to origin function by excluding nucleosomes^3–5^. Despite the occurrence of thousands of ACS in the genome, only 200–300 are efficient for the recruitment of the AAA+ ATPase Origin Recognition Complex (ORC)^6–8^. During the G1-phase, ORC in conjunction with Cdt1 and Cdc6 promotes the binding of the MCM2-7 double hexamer helicase complex giving rise to the pre-replication complex (pre-RC)^9^. The resulting ORC/MCM2-7-bound ARS are said to be licensed for replication initiation and have the ability to initiate replication during the subsequent S-phase^10^.

Replication follows a temporal program of activation during S-phase. ARS are defined by an activation timing based on the observation that some ARS replicate earlier than others^6,8^. Moreover, using DNA combing, it appears that the fraction of cells in a population initiating replication at a given ARS is variable, defining a firing efficiency probability for each ARS^6,7,11^. Nevertheless, timing and efficiency are linked, as inefficient origins tend to fire late during S-phase or to be replicated passively through the use of a neighboring origin^12^. These two interdependent measurements of timing and efficiency are usually used to describe the replication initiation properties of ARS.

In *S. cerevisiae*, many parameters affect ARS activity including limiting transacting factors, different chromosomal location and/or sub-nuclear localization^13^. ARS activity also depends on the chromatin context and histone modifications. First, early origins have a wider NDR than late ARS and adjacent nucleosomes are more precisely positioned^14,15^. Moreover, the strength of ORC recruitment correlates with ARS activity and is itself important for NDR establishment^4,16^.

Second, early ARS activation during S-phase depends on histone acetylation^17,18^. Indeed, the Class I Histone Deacetylase (HDAC) Rpd3 delays initiation of a huge number of replication origins^18–20^ and narrows their nucleosome depleted regions^14^.

Numerous studies attempting to consider transcription as another parameter to define replication initiation led to conflicting results. On one hand, transcription revealed some positive links with replication initiation, as highly transcribed genes were proposed to replicate earlier than low expressed genes^21^. Furthermore, stalled RNA polymerase II (RNAPII) was involved in the recruitment of ORC complex at the rDNA locus, and the activity of many replication origins depends on the presence of specific transcription factor binding sites^22,23^. On the other hand, active ARS are excluded from annotated ORFs and tend to localize after 3’-transcription terminators suggesting that transcription and replication initiation do not coexist^1^. Furthermore, natural or artificial induction of transcription through origins leads to replication defects *via* dissociation or sliding of the pre-RC complex and MCMs respectively^24–28^. In this study, we aimed at clarifying the role of transcription in replication initiation in the context of pervasive transcription.

The analysis of appropriate mutants and the development of new tools to examine nascent transcription have revealed that the RNAPII transcriptional landscape in eukaryotic genomes extends far beyond mRNAs and stable noncoding RNAs^29,30^. While 85% of the *S. cerevisiae* genome is transcribed, only 22% is devoted to protein-coding genes^31^. One source of pervasive transcription stems from widespread initiation at NDRs, an event controlled through early termination by the Nrd1-Nab3-Sen1 (NNS) complex recruited at the 5’ end of all RNAPII transcription units through interaction with RNAPII C-terminal domain (CTD)^32,33^. Recognition by Nrd1/Nab3 of specific motifs on the nascent RNA induces RNAPII termination usually within the first kilobase of transcription in a process coupled to degradation by the nuclear exosome component Rrp6^34^. These cryptic unstable transcripts (CUTs) are revealed in the absence of Rrp6^35–39^. Similarly, depletion of Nrd1 results in transcriptional readthrough and accumulation of Nrd1 unterminated transcripts (NUTs), most of which correspond to extended CUTs^30,40^. Another source of pervasive transcription is linked to mRNA 3’ end formation. The cleavage and polyadenylation (CPF) and cleavage factor (CF) complexes cleave the nascent mRNA just upstream of RNAPII. RNAPII release depends on a CPF-induced allosteric modification of the elongation complex as well as on the digestion of the generated 3’ fragment by the 5’-3’ exonuclease Rat1 (for a recent review see^41^). Thus, before being caught up by the so-called “torpedo”, RNAPII continues transcription leading to an average 160bp-termination window after the polyadenylation site. Inefficient cleavage and polyadenylation can increase the level of this natural source of pervasive transcription^42–44^.

Using nascent transcription and replication analyses in strains depleted for early termination activities, we delineate how pervasive transcription negatively influences replication initiation by remodeling the chromatin structure of ARS. Our study clearly defines pervasive transcription as a new parameter regulating replication initiation.

## RESULTS

### CUTs and NUTs are enriched at early and efficient ARS

To determine the overlap of NUTs and CUTs with replication origins, we defined a list of 234 ARS (Supplementary Table 1) with previously annotated ACS^1,14^ and for which replication timing and efficiency had been established^6^. Among the 234 well-defined ARS used in this analysis, 52 (22%) overlap with a CUT, a NUT or both (Figure 1A, Supplementary Figure 1A)^39,40^. The presence of CUTs and NUTs over 52 replication origins indicates that termination of noncoding transcription is particularly robust around these ARS. Notably, the 52 ARS enriched with NUTs and/or CUTs (ncARS) are replicated earlier and more efficiently than the remaining 182 ARS (Other ARS) (Figure 1B). These observations suggest that non-coding transcription termination may be an important determinant of ARS replication timing and efficiency.

**Figure 1:**
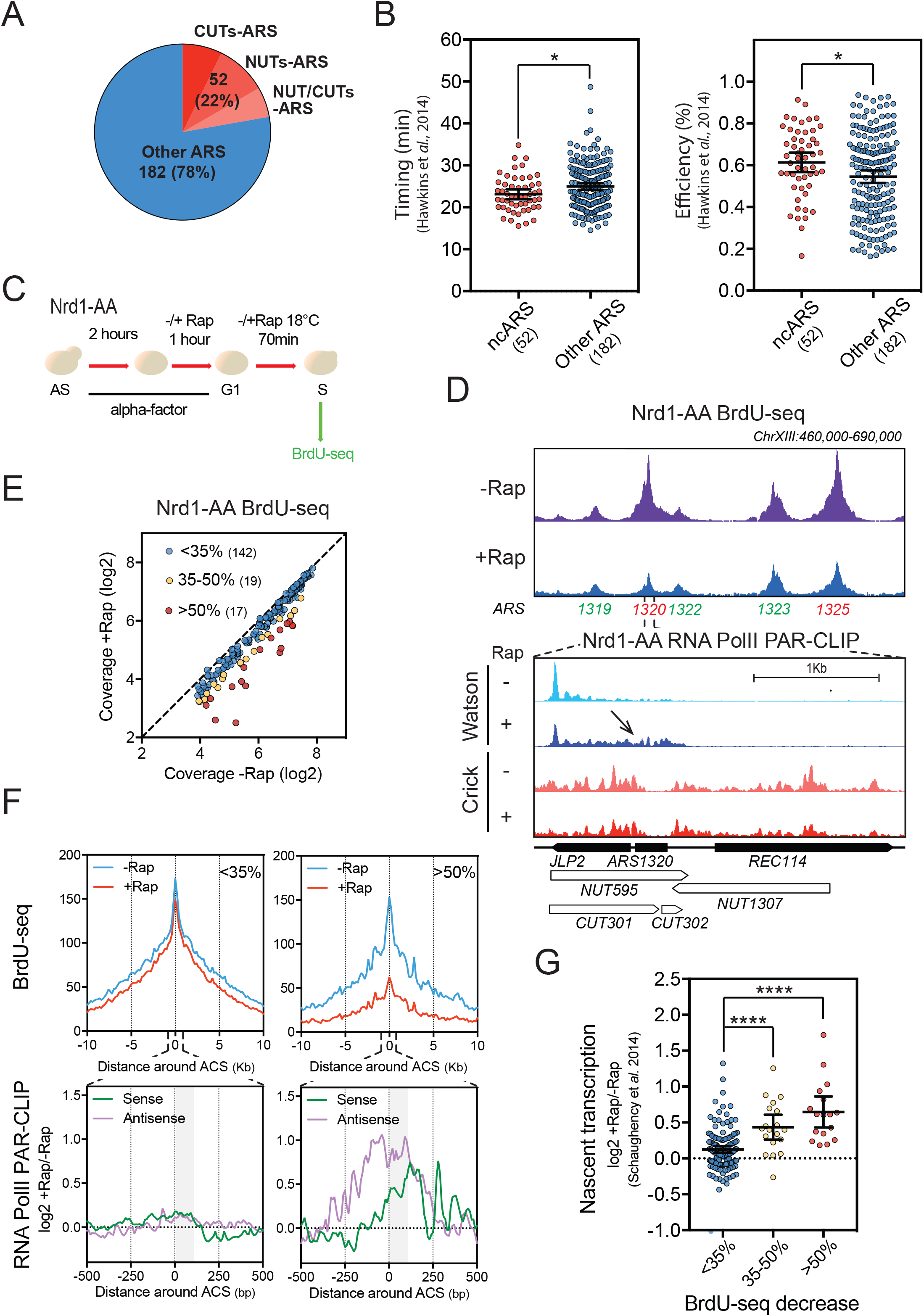
NUTs and CUTs-containing ARS are downregulated when early termination of non-coding RNAs is abrogated. **(A)** Numbers and proportions of ARS overlapping with CUTs, NUTs or CUTs/NUTs. ARS annotations used in this study are listed in Supplementary Table 1. **(B)** Scatter dot plot indicating the timing and efficiency of the noncoding RNA-containing ARS (ncARS) compared to replication origins devoid of overlapping ncRNA (Other ARS). Timing and efficiency data were retrieved from Hawkins et al., 2014. **(C)** Nrd1-AA cells were synchronized in G1-phase with alpha-factor during 3 hours at 30°C. During the last hour, rapamycin (Rap) was added or not in the medium. Cells were then washed and released into the cell cycle at 18°C in the presence of BrdU and −/+Rap. After 70min, cells were collected for DNA extraction and BrdU-seq. **(D)** Snapshot depicting a part of chromosome XIII for the BrdU-seq. Affected ARS are indicated in red and nonaffected in green. Bottom panel shows a zoom around ARS1320 of the RNA PolII PAR-CLIP in the Nrd1-AA strain^30^. Transcriptional readthrough is indicated by an arrow. **(E)** Plot depicting the mean coverage of BrdU nascent DNA in a 5-Kb window around ACS in –Rap versus +Rap. The 17 red dots and 19 yellow dots represent the ARS showing at least 50% and 35–50% decrease in BrdU incorporation in +Rap respectively. Blue dots represent the ARS defined as nonaffected in BrdU incorporation. **(F)** Top: Metagene analysis of the BrdU-seq for the 142 non-affected ARS (<35%) and the 17 most affected ARS (>50%). Profiles represent the mean coverage smoothed by a 200bp-moving window. ARS were oriented according to their ACS T-rich sequence. Bottom: Metagene profiles of the ratio log2 +Rap/-Rap of the RNA PolII PAR-CLIP signal 500bp around the oriented ACS of the least and most affected ARS^30^. Plots were smoothed by a 10bp-moving window. Since ARS are oriented, nascent transcription going towards replication origins is defined as Sense and Antisense. The grey box represents the window in which transcriptional readthrough was analyzed in Figure 1G. **(G)** Scatter dot-plots representing the ratio log2 +Rap/-Rap of the RNA PolII PAR-CLIP signal over the 3 classes of ARS defined in Figure 1E. Total nascent transcription in +Rap and –Rap was defined on oriented ARS by adding the RNA PolII PAR-CLIP mean densities between the ACS to +100bp on the sense strand to the signal over the same region on the antisense strand in each condition using the data from Schaughency et al., 2014. Each 100bp segment was considered as 1 bin (See Materials and Methods).

### Non-coding transcription readthrough affects replication initiation

To investigate the effect of non-coding transcription readthrough on replication, early termination of non-coding RNAs was abrogated by rapid nuclear depletion of Nrd1 through anchor away (AA)^45^. Nrd1 depletion, induced by addition of Rapamycin (Rap) to the engineered Nrd1-AA strain, is accompanied by transcription elongation and accumulation of NUTs^40^. To examine the effect on replication, the Nrd1-AA strain was treated with alpha factor to synchronize the cells in G1-phase and incubated an additional hour −/+ Rap to induce non-coding transcription. Cells were then released from G1 arrest in the presence of BrdU (Figure 1C). FACS analyses indicate a slight cell cycle delay at 80 min in cells depleted for Nrd1, with an increased number of cells in G1 in +Rap compared to –Rap (Supplementary Figure 1B). Samples were harvested for BrdU-seq at 70min after G1-phase release. Visualization of the data revealed a number of well-defined peaks centered on specific ARS (Figure 1D, Supplementary Figures 1C and 1D). Global analysis of the BrdU-seq showed that out of the 178 selected early ARS, 36 (20.2%) present a reproducible, more than 35% decrease in BrdU incorporation in + Rap versus –Rap (17 show >50% decrease and 19 between 35-50% decrease) (Figure 1E). Consistently, metagene analysis from −10Kb to +10Kb around the ACS of the >50% affected ARS revealed a substantial reduction in the BrdU-seq profile in the presence of Rap, while the curves including the <35% affected ARS presented only a slight change in +Rap versus –Rap (Figure 1F). Affected ARS showed a nice overlap with the NUTs-containing ARS defined in Figure 1A (Supplementary Figure 1E). To define whether the decreased BrdU-seq signal of affected ARS upon Nrd1 depletion was linked to transcription, RNAPII PAR-CLIP data from the Corden lab^30^ were used to examine the level of nascent transcription over ARS when depleting Nrd1. Strikingly, the ARS with the strongest decrease in BrdU incorporation in +Rap versus –Rap also showed the highest increase in nascent RNAPII transcription into the ACS to +100bp ORC-footprinting area (Figures 1D and G)^16^. Thus, the replication defect observed following Nrd1 depletion is directly linked to increased nascent transcription through the affected ARS.

RNA and BrdU analyses were also performed with the Rrp6-AA strain. Anchor away of Rrp6 has already been described by our lab to result in CUT elongation^46^. Depletion of Rrp6 resulted in similar effects on ncRNA accumulation and replication initiation (Supplementary Figure 2).

Overall, these data suggest that replication initiation may be hindered by non-coding readthrough transcription.

### High non-coding readthrough transcription leads to ARS chromatin remodeling

Given the links between replication initiation and chromatin structure, we then analyzed the effects of the Nrd1 depletion-induced transcription readthrough into replication origins on nucleosome positioning. To do so, chromatin was extracted from Nrd1-AA cells either untreated or treated for 1 hour with Rapamycin and digested with micrococcal nuclease (MNase). First, sequencing of the 120-200bp fragments protected by nucleosomes revealed the typical NDR around the ACS as previously described (Figure 2A)^4^. Second, analysis of nucleosome positioning at the classes of ARS defined in Figure 1E, revealed a statistically significant increased occupancy in the NDRs of the ARS that are the most affected for replication initiation and present a higher increase in readthrough transcription (Figures 2A, B and C and Figure 1G). This result suggests that high levels of pervasive transcription into replication origins leads to chromatin remodeling which may in turn perturb replication initiation. However, additional parameters are likely to influence replication since the mildly affected ARS (35–50%) do not show significant chromatin remodeling although they are affected in replication initiation.

**Figure 2:**
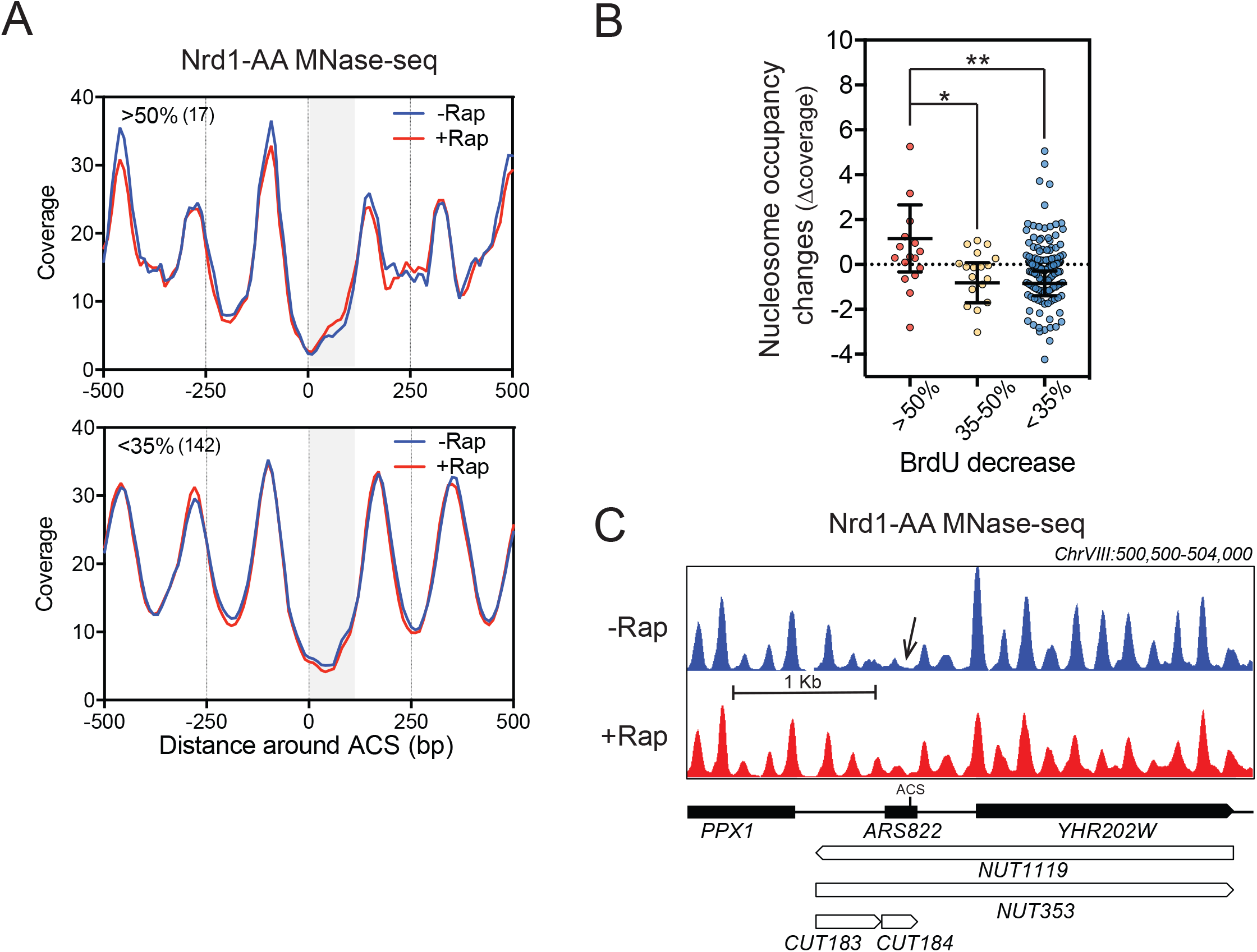
Non-coding transcription readthrough into replication origins alters nucleosome occupancy. **(A)** Metagene analysis of MNase-seq from Nrd1-AA cells treated or not with rapamycin for 1h at the 3 classes of ARS defined in Figure 1. The grey box represents the window in which transcriptional readthrough was analyzed. Only paired-end reads from 120-200bp length were considered. **(B)** Scatter dot plot representing the difference of coverage (Δcoverage) between the ACS to +100 of oriented ARS when comparing +Rap and –Rap conditions. **(C)** Snapshot of the MNase-seq around ARS822 belonging to the class of most affected ARS.

### ARS with high basal level of readthrough transcription are late and inefficient

Recent data show that non-coding transcription occurs all over eukaryotic genomes^35^ and our results indicate that non-coding transcription is detrimental for replication initiation. These observations led to the hypothesis that differences in nascent transcription between ARS may influence both their activity and chromatin structure at steady-state. First, we confirmed that the production of stable transcripts, defined by RNA-seq, strongly drops in the vicinity of the 234 replication origins (including both early and late origins), while profiles of nascent transcription indicate that RNA PolII density stays relatively constant through these ARS (Figures 3A and B). These observations clearly establish that non-coding transcription through replication origins is a frequent event. The set of ARS was then analysed for the level of natural basal readthrough transcription over a 100bp segment between the ACS and the B elements using published RNA PolII PAR-CLIP data^30^. ARS were subdivided in 3 groups according to their natural readthrough transcription levels into this region using a non-biased k-means clustering approach (Figure 3C). The highly transcribed ARS were significantly enriched in ARS lying between two convergent genes (Supplementary Figure 3A). Importantly, nascent transcription towards the ARS measured from upstream and downstream of the oriented ACS were also significantly different for the three groups (Supplementary Figure 3B), indicating that high ARS readthrough mainly stems from higher levels of nascent transcription from the adjacent convergent genes. Interestingly, running these three groups of ARS through replication timing and efficiency data^6^ revealed that the 72 ARS with high readthrough transcription have a significantly delayed replication timing and reduced replication efficiency compared to the ARS with lower transcriptional readthrough (Figure 3D). Notably, the 52 NUTs/CUTs-containing ARS present significantly lower natural readthrough transcription than the highly transcribed ARS and appear more similar to the mildly transcribed ARS, further supporting that NNS-mediated termination positively contributes to replication timing and efficiency by shielding ARS form pervasive transcription (Supplementary Figure 3C). Consistent with these results, earlier defined ORC-bound ARS (ORC-ARS) exhibit significantly less readthrough transcription compared to non ORC-bound and non-replicated ARS (nr-ARS) (Supplementary Figure 3D)^4^.

**Figure 3:**
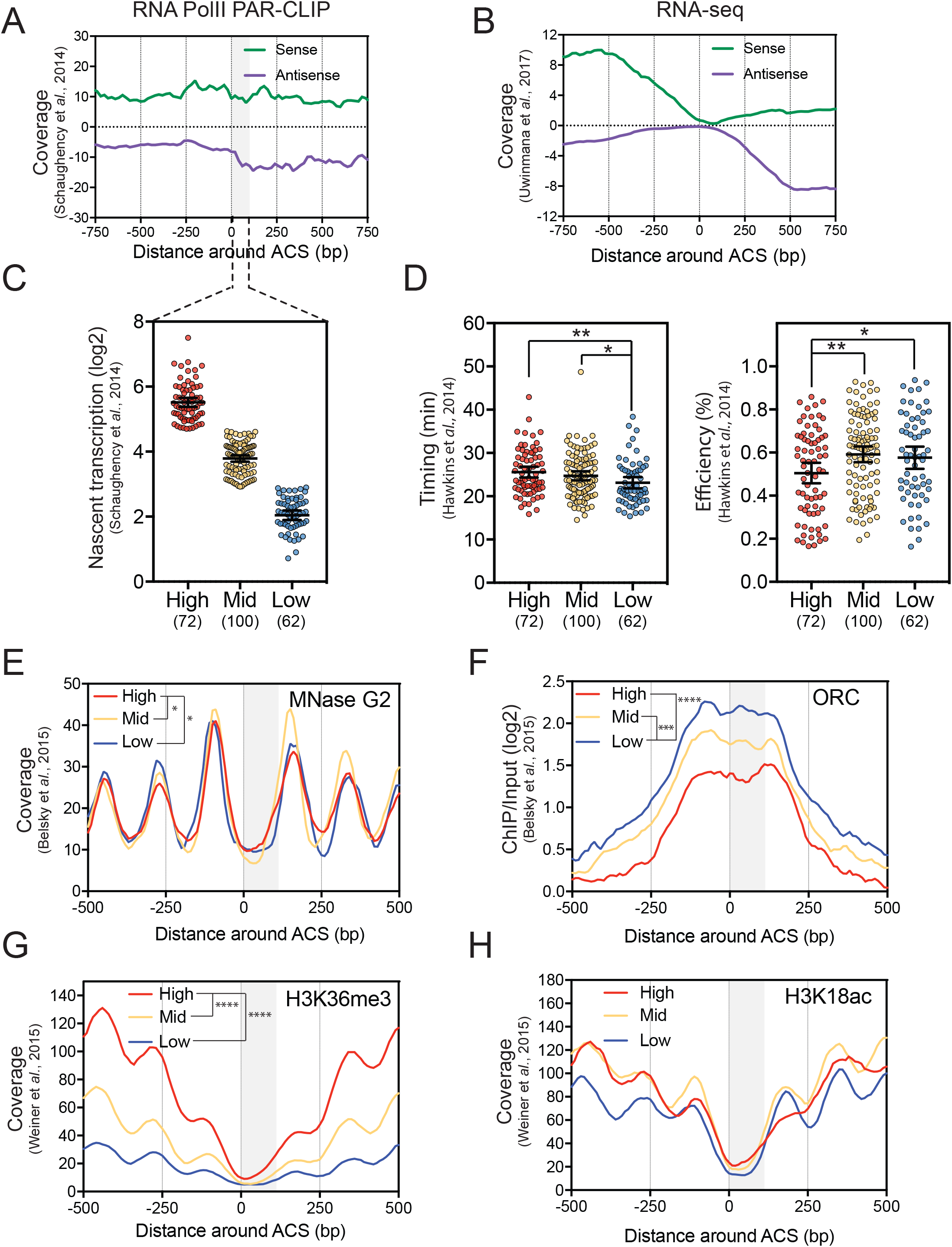
Natural pervasive transcription correlates with timing/efficiency and specific chromatin features of replication origins. **(A)** Metagene analysis of Sense and Antisense RNA PolII PAR-CLIP^30^ and **(B)** RNA-seq^75^ data in the vicinity of 234 oriented replication origins. Data were smoothed by a 20bp-moving window. The grey box represents the window in which transcriptional readthrough was analyzed. **(C)** The 234 replication origins were divided into 3 classes (High, Mid or Low) according to their level of total natural readthrough transcription using a non-biased k-means clustering approach. This basal nascent transcription was calculated on oriented ARS by adding the RNA PolII PAR-CLIP mean densities between the ACS and +100bp on the sense strand to the signal over the same region on the antisense strand. Each 100bp segment was considered as 1 bin. Natural nascent transcription data were taken from Schaughency et al., 2014. **(D)** Replication timing and efficiency of the 3 classes of ARS defined as in (C). **(E) (F) (G) (H)** Nucleosome positioning, ORC, H3K36me3, and H3K18ac levels considering the 3 classes of replication origins. Data for nucleosome positioning, ORC recruitment and histone marks were retrieved from Belsky et al., 2016 and Weiner et al., 2015 respectively. For these plots, ARS were oriented and aligned according to their ACS T-rich sequence. The significance of differences between the 3 classes was calculated over the ACS to +100bp region considered as 1 bin.

Thus, natural pervasive transcription into the ORC-binding area appears to be anti-correlated with ARS function as defined by its timing and efficiency as well as its ability to bind the ORC complex.

### Steady-state highly transcribed ARS present distinctive chromatin features

Using recently published nucleosome occupancy data^47,48^, the 72 ARS with high pervasive transcription levels appear to be associated with significant increased nucleosome occupancy in G2-phase over the ORC-binding area compared to the 162 with lower transcription levels (Figure 3E). Such a significant correlation is not detected in G1-phase, possibly due to components of the positioned pre-RC complex that may lead to the protection of non-nucleosome 120–200bp fragments^15,16^ (Supplementary Figure 4A). Thus, high levels of nascent transcription into the ACS to +100bp area correlate with higher nucleosome occupancy when the pre-RC is not assembled. Accordingly, low levels of pervasive transcription are associated with higher ORC binding (Figure 3F).

We also analyzed the correlations between natural pervasive transcription into ARS and histone modifications^48^. We found that H3K36me3 levels over the ACS-100bp area positively correlate with the levels of nascent transcription (Figure 3G). We did not detect a significant correlation between nascent transcription and H3K18, H3K14, H4K12 and H4K5 acetylation over the ORC-binding region (Figure 3H and Supplementary Figures 4B, C and D). However, we detected a significant lack of these acetylation marks at the downstream nucleosome of the highly transcribed class (Supplementary Figure 4F).

Interestingly, H3K36me3 is deposited by the Set2 histone methyl transferase and leads to the recruitment of the Rpd3 HDAC known to deacetylate the lysines cited above and described as being involved in replication control^19,49^. In contrast, the two groups of ARS present no difference in H3K4me2, another mark promoting deacetylation of the chromatin via binding of the Set3 HDAC (Supplementary Figure 4E)^50^.

Together these observations support the view that pervasive transcription through an ARS promotes the formation of a closed chromatin structure reducing its ability to interact with ORC, thereby decreasing its efficiency and/or delaying its replication timing.

### Non-coding and mRNA readthrough transcription over ARS induce H3K36 methylation, histone deacetylation and increased nucleosome occupancy

To strengthen the causal relationship between nascent transcription, changes in chromatin organization and ARS activity, we examined the effect of induced transcription on ARS chromatin remodeling. We first ranked the replication origins according to their increase in transcriptional readthrough levels in the Nrd1-AA strain and to their decrease in BrdU incorporation during early S-phase. We picked three ARS belonging to the most affected ARS in BrdU incorporation and presenting high levels of induced transcriptional readthrough (Figure 4A). As a control, we took two replication origins belonging to the least affected ARS in replication and showing no or weak induced readthrough. To relate the replication defect induced by non-coding readthrough transcription to changes in chromatin structure, chromatin immunoprecipitation was used to compare Histone H3 occupancy, H3K36 methylation and H3K18 acetylation in −/+ Rap. Primers for qPCR were designed to target the NDR of the ARS. At the 3 affected ARS, rapamycin treatment resulted in increased H3 occupancy and H3K36 methylation as well as a decrease in H3K18 acetylation, while no changes were observed at the non-affected ARS (Figures 4B).

**Figure 4:**
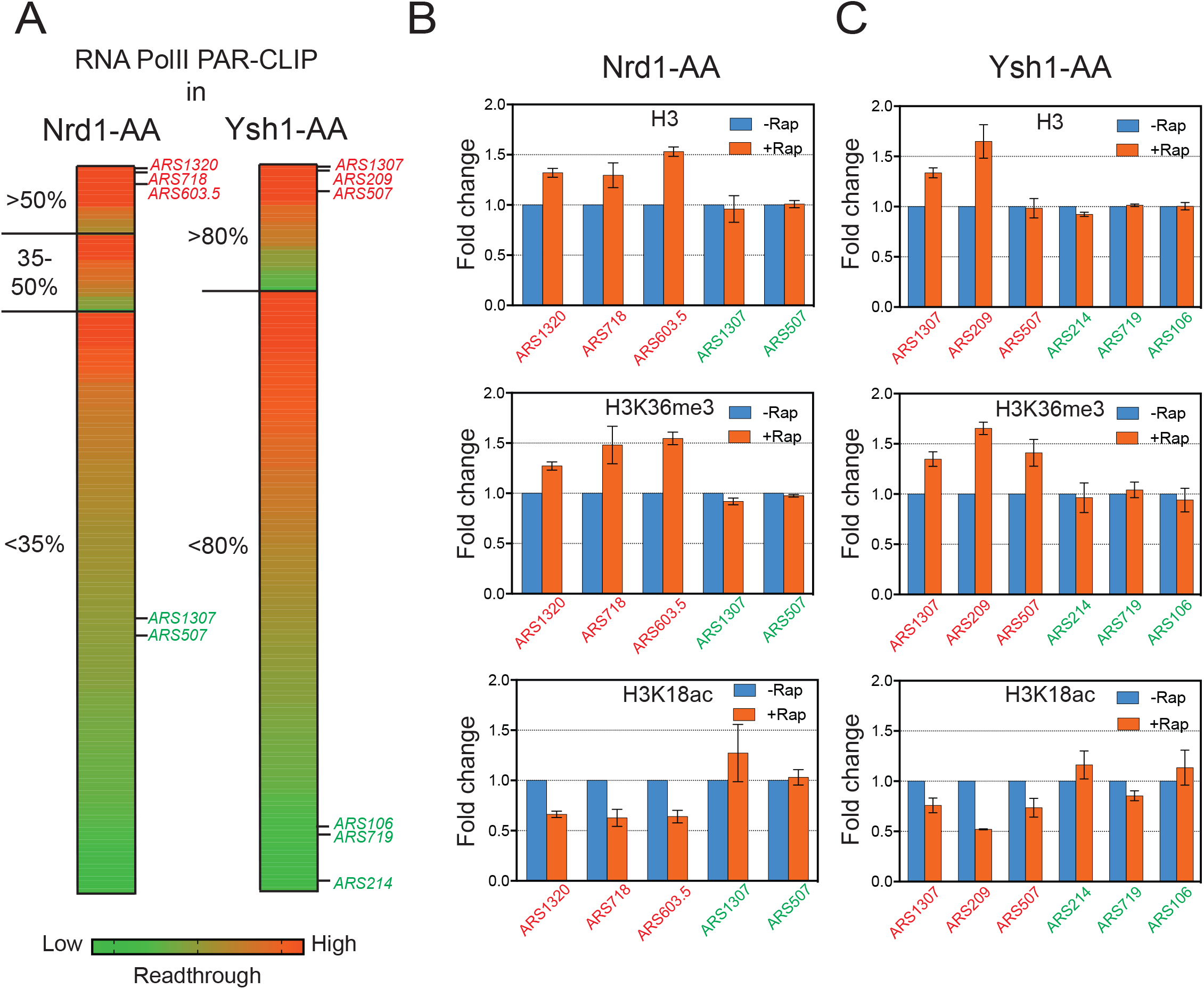
Non-coding and mRNA transcription readthrough at replication origins leads to chromatin remodeling. **(A)** Heat map representing the fold change of transcriptional readthrough at replication origins in Nrd1-AA and Ysh1-AA mutants. The 178 ARS were classified according to their BrdU incorporation defects and ranked by their total readthrough increase on the ACS to +100bp. The 3 ARS indicated in red for each mutant have high levels of total readthrough. The ARS depicted in green show mid or low total readthrough and have been used as controls for the following experiments. **(B)(C)** Chromatin immunoprecipitation (ChIP) of H3, H3K36me3, H3K18ac at ARS with high (red) and low (green) readthrough transcription. Asynchronous cells were treated 1h or 30min with rapamycin to induce Nrd1 and Ysh1 depletion from the nucleus respectively. ChIP was performed as described in Materials and Methods. Immunoprecipitated ARS loci were normalized to immunoprecipitated *SPT15* ORF after qPCR amplification. Fold enrichment was artificially set to 1 for the –Rap condition (n=3). Error bars represent Standard Error of the Mean (SEM).

Since 86% of replication origins are located in the vicinity of a convergent coding gene, we decided to anchor away the essential mRNA 3’ cleavage and polyadenylation factor (CPF/CF) endonuclease Ysh1. As expected, nuclear depletion of Ysh1 has a major impact on replication progression and more specifically on replication initiation, as most of the 178 considered ARS showed reduced BrdU incorporation in the presence of rapamycin, with 31 ARS presenting more than 80% decrease (Supplementary Figures 5A, B, C and D). Combining BrdU-Seq in Ysh1-AA −/+ Rap with published RNA PolII PAR-CLIP data of this strain^30^ revealed that the 31 ARS with the strongest replication defect also present the highest increase in readthrough transcription (Supplementary Figures 5D and E). Replication origins were then ranked according to their induced levels of readthrough and their defects in early replication as performed for the Nrd1-AA strain. Chromatin immunoprecipitation revealed increased H3 occupancy for the ARS showing high levels of mRNA readthrough (with the exception of ARS507) while H3K36me3 levels increased and H3K18ac levels decreased.

Taken together, these experiments demonstrate that increased noncoding and mRNA readthrough transcription causes increased nucleosome occupancy and histone deacetylation at the downstream ARS, two parameters described to interfere with the efficiency of ORC binding and ARS licensing^14,18–20^.

### Replication defects induced by non-coding readthrough transcription are partially rescued in the absence of H3K36 methylation

To decipher the molecular cascade of events, we analyzed the effects of non-coding transcription readthrough on replication initiation in a Nrd1-AA *set2Δ* strain (Figure 5A). Global analysis of replication by flow cytometry indicates that replication is still delayed in the absence of H3K36 methylation in the Nrd1-AA background (Supplementary Figure 6A). However, by taking the same classes of defective ARS as defined in Figure 1, we observed a partial rescue of BrdU incorporation (Figures 5B, C and D) although non-coding RNAs were still produced in the absence of Set2 (Supplementary Figure 6B). These observations support the view that pervasive transcription drives chromatin remodeling of replication origins, which in turn defines, at least in part, ARS activity.

**Figure 5:**
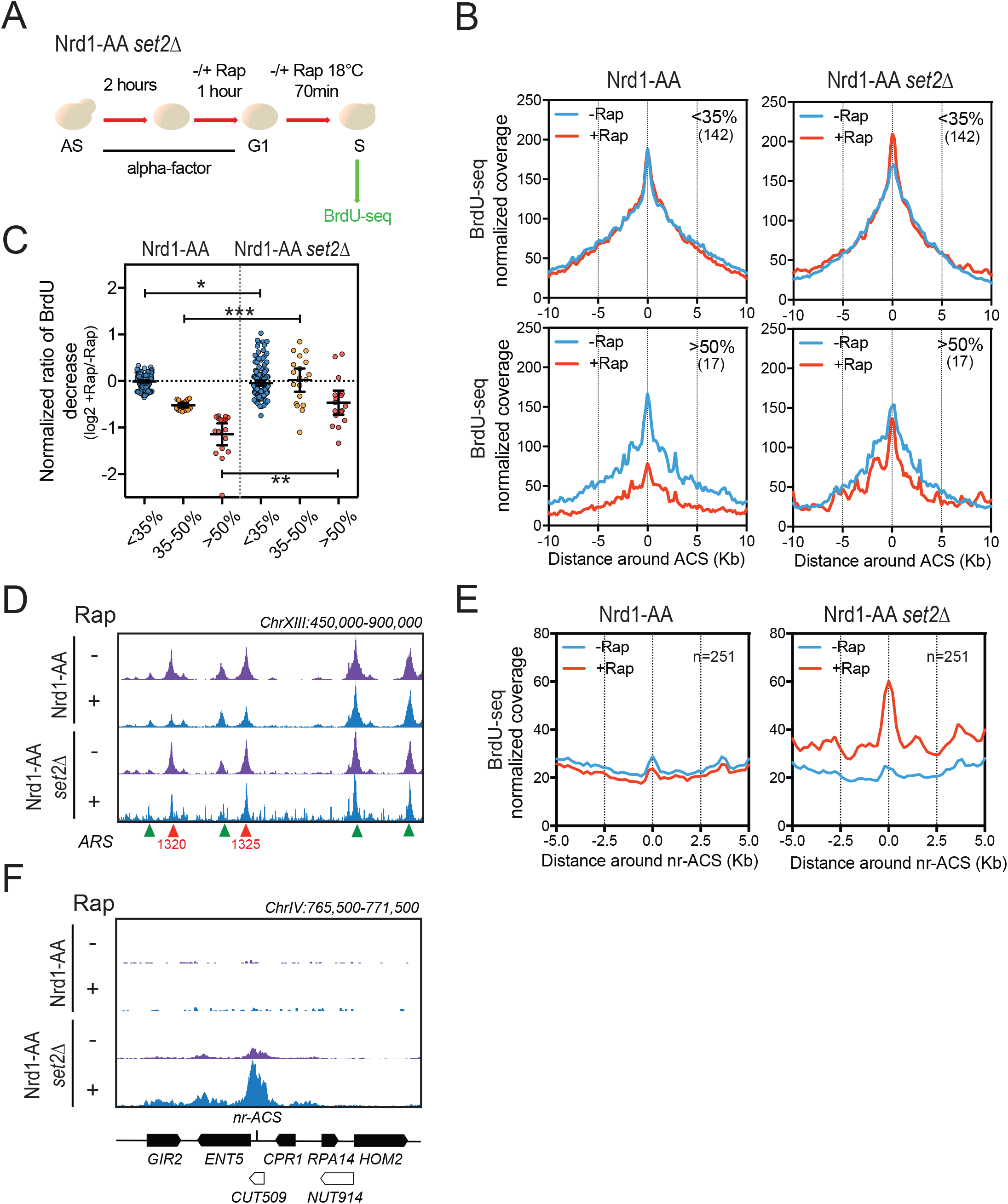
Absence of Set2 H3K36 methyl transferase partially rescues replication defects due to non-coding transcription readthrough. **(A)** Experimental scheme as described in Figure 1C. **(B)** Metagene analysis of the BrdU-seq for the 142 non-affected ARS (<35%) and the 17 most affected ARS (>50%) for the Nrd1-AA and Nrd1-AA *set2Δ* strains. Plots for the Nrd1-AA strain were already presented in Figure 1F with the exception of the Normalized coverage for which calculation is detailed in Materials and Methods. Profiles represent the mean coverage smoothed by a 200bp-moving window. ARS were oriented according to their ACS T-rich sequence. **(C)** Scatter dot plot presenting the normalized BrdU ratio for the different classes of ARS affected in BrdU incorporation in an Nrd1-AA strain and for the same classes of ARS in the Nrd1-AA *set2Δ* strain. **(D)** Snapshot depicting the BrdU-seq reads for a part of chromosome XIII. ARS that are rescued in BrdU incorporation in the Nrd1-AA *set2Δ* +Rap condition are depicted in red while non-affected ARS are in green. **(E)** Metagene analysis of BrdU incorporation 5Kb around non-replicating ACS (nr-ACS) in the indicated strains grown in −/+Rap. The representation is smoothed over a 200bp-moving window. **(F)** Snapshot illustrating the activation of a dormant nr-ACS in the Nrd1-AA *set2Δ* +Rap condition.

Interestingly, rapamycin treatment of the Nrd1-AA *set2Δ* strain led to the appearance of small BrdU incorporation peaks all over the genome (Figure 5D), suggesting that replication initiation loses its specificity when both non-coding transcription termination and H3K36 methylation are abrogated. Moreover, analysis of BrdU incorporation around non-ORC bound and non-replicated ACS^4^ revealed an increase of replication initiation at non-canonical sites when noncoding transcription readthrough is induced in the absence of Set2 (Figures 5E and F). Thus, while non-coding transcription interferes with replication in the presence of Set2 by favoring a closed chromatin structure, transcription over dormant ARS in the absence of Set2 activates replication, probably as a result of nucleosome instability.

## DISCUSSION

We have shown that ncRNA early termination by the Nrd1-dependent pathway in the vicinity of a subset of ARS protects these origins from transcription and replication initiation defects. These observations suggest that inefficient non-coding transcription termination may influence replication origin activity. Consistently, our analyses reveal that natural readthrough transcription correlates with reduced ARS activity and a specific replication origin chromatin structure. Moreover, using mRNA or cryptic transcription termination mutants, we have established that nascent transcription is the causal link defining chromatin organization at a number of ARS. Finally, we have presented evidence that pervasive transcription-induced chromatin modifications and not only pervasive transcription *per se* controls ARS activity. Thus, we propose pervasive transcription as a novel primordial parameter defining replication initiation features (Figure 6).

**Figure 6:**
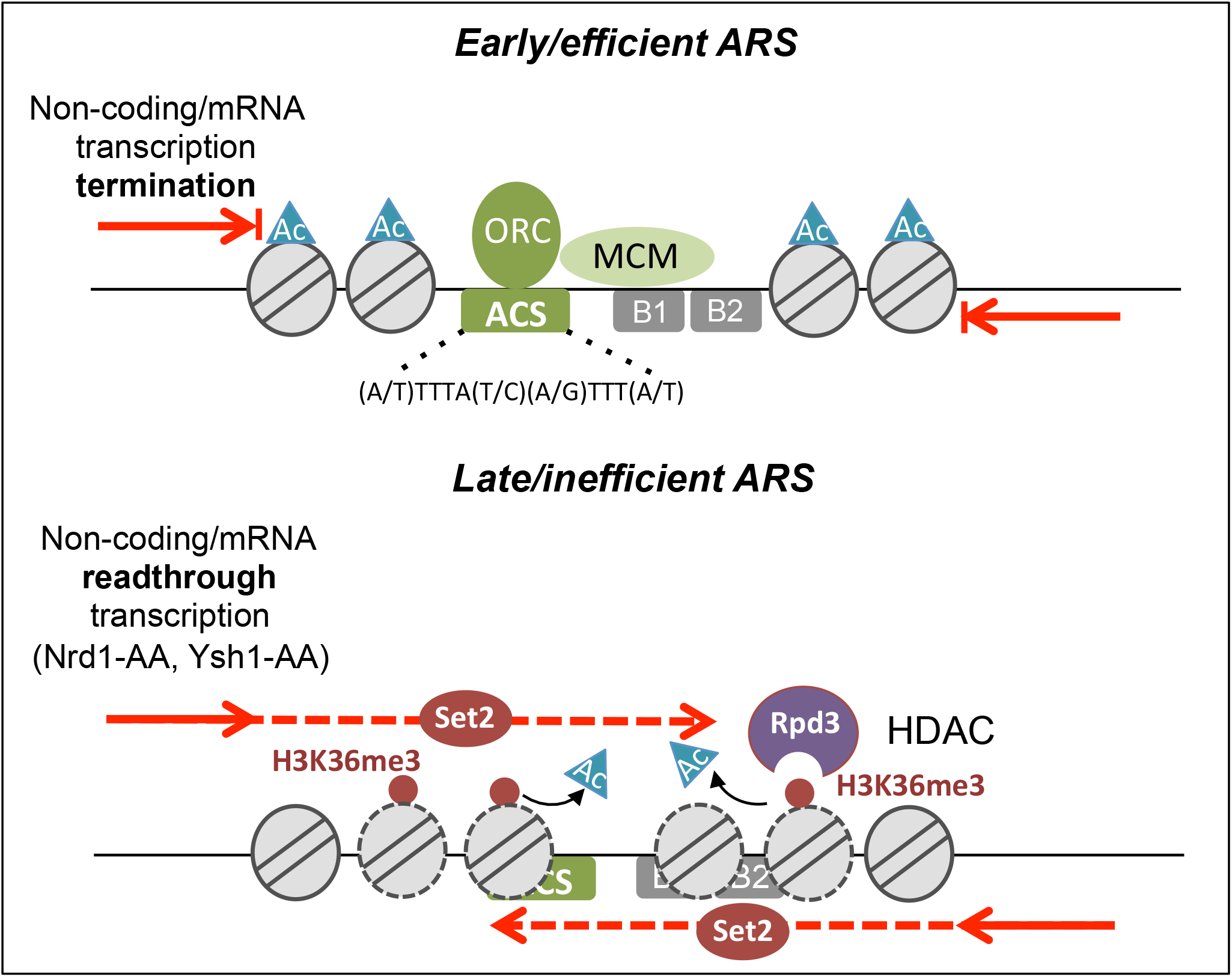
Pervasive transcription influences replication timing/efficiency by modulating ARS chromatin structure and ORC binding. The chromatin structure of replication origins is defined, at least in part, by the level of pervasive readthrough transcription. In the presence of efficient noncoding (Nrd1/Nab3/Sen1-dependent) or mRNA (CPF/CF-dependent) transcription termination, ARS present low H3K36 trimethylation, high downstream nucleosome acetylation (Ac), a wide NDR and more ORC binding at the ACS, favoring early and efficient replication. If transcription termination is deficient, H3K36me3 by Set2 increases and histone acetylation decreases likely through the recruitment of the Rpd3 histone deacetylase; these modifications increase nucleosome stability and occupancy over the ARS, lowering the level of ORC recruitment and resulting in late and inefficient ARS replication.

### Nrd1-dependent transcription termination protects a subset of early/efficient replication origins from pervasive transcription

Previous studies on the relationship between transcription and replication have led to conflicting observations. Our analyses of pervasive transcription lead to the conclusion that transcriptional readthrough at ARS is detrimental for replication initiation as already proposed by some reports^25–27,51^. Importantly, however, we show that natural nascent transcription *per se* is a criterion defining ARS activity genome-wide. Thus, the strategy for a replication origin to increase its activity would be to limit pervasive transcription. Accordingly, a subset of early and efficient ARS are protected from pervasive transcription thanks to surrounding non-coding transcription termination by the Nrd1-dependent pathway (Figures 1, 3 and Supp. Figure 3C). Thus, we propose that Nrd1-dependent termination in the vicinity of a replication origin is an efficient way to decrease transcriptional readthrough. Since Nrd1-dependent termination is regulated under stress conditions, it is tempting to speculate that this might also impact on replication origins usage^52,53^. This scenario would define a novel role for non-coding RNAs in the regulation of genome maintenance.

Interestingly, no mechanism similar to Nrd1-dependent termination has been described in other eukaryotes yet^54^. In *S. cerevisiae*, Nrd1-protected origins reach a median firing efficiency peaking at 58%, while firing efficiency is around 30% in *Schizosaccharomyces pombe*^55^. An attractive view is that Nrd1-dependent transcription termination represents an evolutionary pathway maximizing replication initiation efficiency.

Importantly, there is no significant linear correlation between nascent transcription and ARS activity. This connection appears when ARS are divided into subsets (Figure 3). These observations indicate that nascent transcription influences ARS activity only beyond a certain threshold and that other parameters contribute to origin function. Accordingly, a variety of molecular events have been involved in regulating ARS activity, which include MCM levels bound to ARS, cell cycle regulated binding and affinity of the ORC complex for the ACS, or the presence of Fkh1/2 proteins^16,56–59^.

### Replication origin chromatin structure is influenced by pervasive readthrough transcription

Previous work has involved the chromatin structure at replication origins as a parameter defining replication initiation. Early ARS tend to show an open chromatin, low H3K36me3 and high histone acetylation levels^14,60^. Our results indicate that these features may be directly related to the level of natural nascent transcription. Indeed, when compared to highly transcribed ARS, origins with low level of readthrough transcription are characterized by lower nucleosome occupancy, lower H3K36me3 and higher histone acetylation levels (Figure 3). Notably, H3K36me3 is deposited by Set2, a histone methyl transferase (HMT) recruited through interaction with the C-terminal domain (CTD) of the elongating RNAPII. H3K36me3 serves as a platform for the binding of Rpd3S, a histone deacetylase complex described to deacetylate and stabilize reassembled nucleosomes in the wake of the transcription machinery, suppressing initiation from cryptic sites within ORFs^61^ and see^50^ for a recent review. This molecular mechanism may represent the connection between pervasive transcription and chromatin structure of replication origins. Indeed, increasing transcriptional readthrough into replication origins is accompanied by higher levels of H3K36me3, lower levels of H3K18ac, increased nucleosome occupancy and replication defects (Figures 1, 2, 4, and Supplementary Figure 4). Importantly, BrdU-seq experiments cannot discriminate between timing and efficiency defects. However, since nucleosome methylations are relatively stable modifications, it would be appealing to propose that even rare events of pervasive transcription may stably inactivate replication origins until the subsequent S-phase dilutes or a histone demethylase erases these methylation marks. In such a model, pervasive transcription may shape replication origin chromatin for inefficient usage.

These observations shed new light on earlier published results. First, loss of Rpd3 leads to a global increase in acetylation around ARS and early firing of many late origins^18,19^. It was proposed that the “accelerated” replication in *rpd3Δ* is mainly due to a reduced titration of replication initiation factors by the rDNA origins^62^; however, loss of Rpd3 also has a global impact on increasing the size of replication origins NDR^14^. Of note, loss of Rpd3 abrogates the function of both Rpd3S and Rpd3L, another histone deacetylase complex involved in gene repression following recruitment to promoters via a Set2/H3K36met3 independent pathway^50^. While loss of Rpd3L subunits was shown to result in increased BrdU incorporation at a number of ARS, especially those adjacent to upregulated genes, loss of Rpd3S subunits or Set2 led to a weaker but more generalized increase in ARS replication initiation^19^. This difference could reflect a primary specific effect of loss of Rpd3L on rDNA replication and increased availability of replication factors in the absence of this HDAC. Although the replication phenotype was less pronounced when deleting Rpd3S, the data support the view that pervasive transcription promotes Set2/H3K36me3-mediated histone deacetylation by Rpd3S and globally contributes to negatively regulate replication origin activity.

Our results further indicate that high readthrough transcription correlates with decreased ORC binding (Figure 3 and Supplementary Fig. 3D). It was proposed that replication initiation timing depends more on the surrounding chromatin than on the ORC-ACS *in vitro* affinity by itself^56^. It would be interesting to further dissect the molecular events at origins and to define whether nascent transcription *per se* evicts ORC complex, leading to nucleosome deposition and histone modifications, or whether RNAPII readthrough and nucleosome incorporation outcompete ORC turnover (Figure 6).

Surprisingly, we observe genome-wide appearance of BrdU peaks when *SET2* is deleted in conjunction with the anchor-away of Nrd1. These peaks are not detected in the *set2Δ* mutant alone indicating that activation of dormant origins is not only related to the absence of the histone mark. Deletion of *SET2* is known to drastically increase the level of intragenic transcription initiation^63^. A fraction of this novel nascent transcription may be cleared by the Nrd1-dependent termination pathway. We propose that in the absence of Nrd1, transcription may hit dormant origins, which in the absence of H3K36 methylation will promote replication initiation due to nucleosome instability and chromatin opening. Thus, when not associated with H3K36 methylation, transcription may have a positive effect on replication initiation.

### Implications for a pervasive transcription-dependent replication initiation model in metazoans

Identification of replication origins in mouse embryonic stem cells showed that nearly half of them are contained in promoters^64^, and recent ORC binding data in human cells led to the same conclusion^65,66^. In contrast, replication initiation in *S. cerevisiae* more frequently occurs next to gene terminators^1^. Interestingly, human promoters are bidirectional and lead to the production of highly unstable promoter upstream transcripts (PROMPTs), suggesting that pervasive transcription could also play a role in metazoan replication initiation^67–69^. It has recently been shown that replication initiation in human cells occurs within broad over 30kb regions, flanked by ORC binding at one of the two ends^70^. Considering that MCM helicases can slide along the chromosome with the help of transcription^27^, it would be appealing to propose that promoter-associated pervasive transcription redistributes the MCM helicases from their ORC binding initial site of loading. Since we have established pervasive transcription as a novel primordial parameter regulating replication initiation in *S.!cerevisiae*, its importance for the metazoan replication program will deserve being studied in the next few years.

## EXPERIMENTAL PROCEDURES

### Yeast strains and Microbiological methods

All strains were derived from W303 and Anchor-Away genetic backgrounds (see Supplementary Table 2)^45^. Cells were grown in YEPD medium (1% yeast extract, 1% peptone) supplemented with 2% glucose as the carbon source. The BrdU incorporation cassette was introduced according to^71^. Anchor-away of Nrd1-AA, Nrd1-AA *set2Δ*, Rrp6-AA or Ysh1-AA strains was induced by adding 1μg/ml of rapamycin to the medium.

### Cell culture and FACS analyses

Cells grown at 30°C to an OD600= 0.4 were synchronized in G1-phase with α-factor (20 ng/ml, Sigma) for 3h in total. After two washes with distilled water, cells were released into S-phase at 18°C. Depending on the experiment, cells were released in YEPD medium containing BrdU (100 μg/ml, Sigma) and treated or not with rapamycin. Cells were then collected at different times after G1-release depending on the experiment and treated with 0.1% Sodium Azide (Sigma). Flow cytometry was performed on ethanol-fixed cells using propidium iodide (Sigma). Flow cytometry profiles were obtained using Gallios flow cytometer (Beckman-Coulter) before data analyses using FlowJo software (LLC).

### RNA extraction and Reverse Transcription-qPCR

RNAs were extracted using Trizol (Invitrogen). Purified nucleic acids were first treated with DNAse (Ambion) before reverse transcription using SuperScript II (Invitrogen). cDNA was then amplified as described below using the primers available upon request.

### BrdU immunoprecipitation and sequencing

BrdU immunoprecipitation was mainly performed as described in^72^ with the following modifications. Genomic DNA was sonicated into 300-400 bp fragments and denatured. 5μg BrdU antibody (BD PharMingen, 555627) coupled to 75μl of Dynabeads Protein G (Invitrogen) were added to 5μg of denatured BrdU-containing genomic DNA in BrdU IP buffer (PBS + 0.0625% Triton X-100). After 1 hour incubation at room temperature on a wheel, beads were washed twice with BrdU IP buffer and eluted in Tris-HCl 10mM (pH 8.0), EDTA 1mM, 1% SDS. Eluates were cleared of SDS using the NucleoSpin Gel and PCR Clean-up (Macherey-Nagel). Libraries were constructed using the iDeal library preparation kit (Diagenode). Sequencing was performed on the HiSeq 4000 sequencer (Illumina). For specific loci analyses by quantitative PCR, BrdU-containing DNA was amplified using the SYBR select master mix for CFX (Applied Biosciences) on a CFX96 Real-time detection system (Bio-Rad).

### Mnase-seq

MNase treatment was performed as described in^48^. Chromatin was extracted by breaking cells with bead beating in a magnalyser (Roche). Chromatin was then collected by centrifugation and re-suspended in NP-buffer (0.5 mM spermidine, 1 mM β-ME, 0.075% NP40, 50 mM NaCl, 10 mM Tris pH 7.4, 5 mM MgCl2, 1 mM CaCl2). MNase (Thermo-Scientific) treatment was performed at a previously optimized concentration to have comparable intensity of both mono- and di-nucleosomes within and between the samples. MNase treatment was followed by de-crosslinking, protease treatment and DNA was extracted using NucleoSpin gel and PCR extraction columns (Macherey-Nagel). iDeal library preparation kit (Diagenode) was then used for library construction. Sequencing was performed on the HiSeq 4000 sequencer (Illumina).

### BrdU-seq and MNase-seq bioinformatic analyses

50bp paired-end reads were aligned to sacCer3 genome assembly using HTSstation^73^. PCR duplicates were removed from the analysis. For the MNase-seq, 120-200bp fragments were filtered to detect molecules with nucleosome size using HTS Bioscript^73^. BrdU-seq and MNase-seq density files (bigwig) were averaged for the two replicates of each condition. All subsequent analyses were performed using HTS Bioscript including metagene analyses. To assign one value of BrdU incorporation to each ARS, BrdU incorporation was measured 5kb around ACSs considering this area as 1bin. For the MNase-seq, nucleosome occupancy was quantified over the ACS to 100bp area of oriented ARS.

Since no spike-in was used in our experiments and since Nrd1-AA *set2Δ* + Rap substantially changes the density profile because of dormant origins firing, Figure 5 was normalized as follows: the <35% affected ARS class for BrdU incorporation was considered as equal in +Rap and –Rap in average 5kb around the ACS in both Nrd1-AA and Nrd1-AA *set2Δ*. This gave a normalization factor for each strain, which was then used to quantify the other classes.

### Transcriptional readthrough calculation for RNAPII PAR-CLIP

Induced transcriptional readthrough was calculated as follows. First, mean densities of Watson and Crick strands in each condition were calculated on oriented ARS between the ACS to +100bp considering this region as 1bin using HTS bioscript. A pseudo-count of 1 was added to each value to correct for low values, which could lead to overestimated ratios defined hereafter. The total readthrough was then calculated by adding the values obtained for Watson and Crick in +Rap divided by the sum of Watson and Crick in –Rap. For Figure 3, Watson and Crick strand values were added to get the total amount of natural readthrough. Three classes of ARS with different levels of natural readthrough (High,Mid and Low) were then defined through the use of k-means clustering (http://scistatcalc.blogspot.ch/2014/01/k-means-clustering-calculator.html).

### Chromatin Immunoprecipitation (ChIP)

ChIP experiments were performed as described previously with some modifications^74^. Antibodies against H3 (Abcam ab1791), H3K36me3 (Abcam ab9050) and H3K18ac (Abcam ab1191) were incubated with Protein G dynabeads (Invitrogen) before being mixed with sonicated chromatin and incubated on a wheel at 4°C for 2 hours. After washes, immunoprecipitated chromatin was eluted before being purified on columns (Macherey-Nagel). ChIPs were repeated three times with different chromatin extracts from independent cultures. Immunoprecipitated DNA was then purified and quantified by qPCR.

Immunoprecipitated ARS loci were normalized to immunoprecipitated *SPT15* ORF after qPCR amplification.

### List of non-coding RNAs and replication origins

The list of CUTs was obtained from^39^, while the list of NUTs was kindly provided by the Cramer lab^40^. Among the NUTs, only those showing at least a 2-fold increase in +Rap/-Rap were taken into account to unify the threshold of ncRNA definition between CUTs and NUTs. The list of ARS (Supplementary Table 1) consists of the 234 ACS taken from^14^ that overlap with the replication origins described in^6^, for which replication timing and efficiency have been defined. Replication origins with an efficiency <15% were not taken into account.

### Statistical analysis

All statistical analyses of this work were performed using Prism 7.0 (Graphpad). All tests are non-paired tests (with the exception of Figure 5C). T-tests or Mann-Whitney tests were used according to the normality of the data analyzed, which was calculated using a d’Agostino-Pearson omnibus normality test. * for p-value<0.05; ** <0.01; ***<0.001 and ****<0.0001.

### Bioinformatics data availability

The accession number for the raw and processed sequencing data reported in this paper is GEO: GSE111058.

### Published datasets used for analyses

For RNA PolII PAR-CLIP, RNA-seq, chromatin and ORC profiles, data were retrieved from^30^ (GEO: GSE56435),^75^ (GEO: GSE89601),^48^ (GEO: GSE61888) and^16^ (SRA: SRP041314).

### Author Contributions

J.S. and F.S. conceived the study. J.S. performed most experiments and analyzed the results together with F.S.; J.K. performed the MNAse-seq; J.S. and F.S. wrote the manuscript.

## AKNOWLEDGEMENTS

We would like to thank O. Aparicio, S. Bell, D. MacAlpine and P. Cramer for sharing data and reagents; M. Strubin, F. Steiner, M. Shyian, G. Canal, T. Halazonetis and all members from the Stutz lab for comments on the manuscript. We also would like to thank Domenico Libri for communication of unpublished results. This work was funded by the Swiss National Science Foundation (31003A 153331), iGE3 and the Canton of Geneva, as well as a Polish Swiss Research Programme (PSRP NoCore 183/2010).

---

**Supplementary Figure 1:**
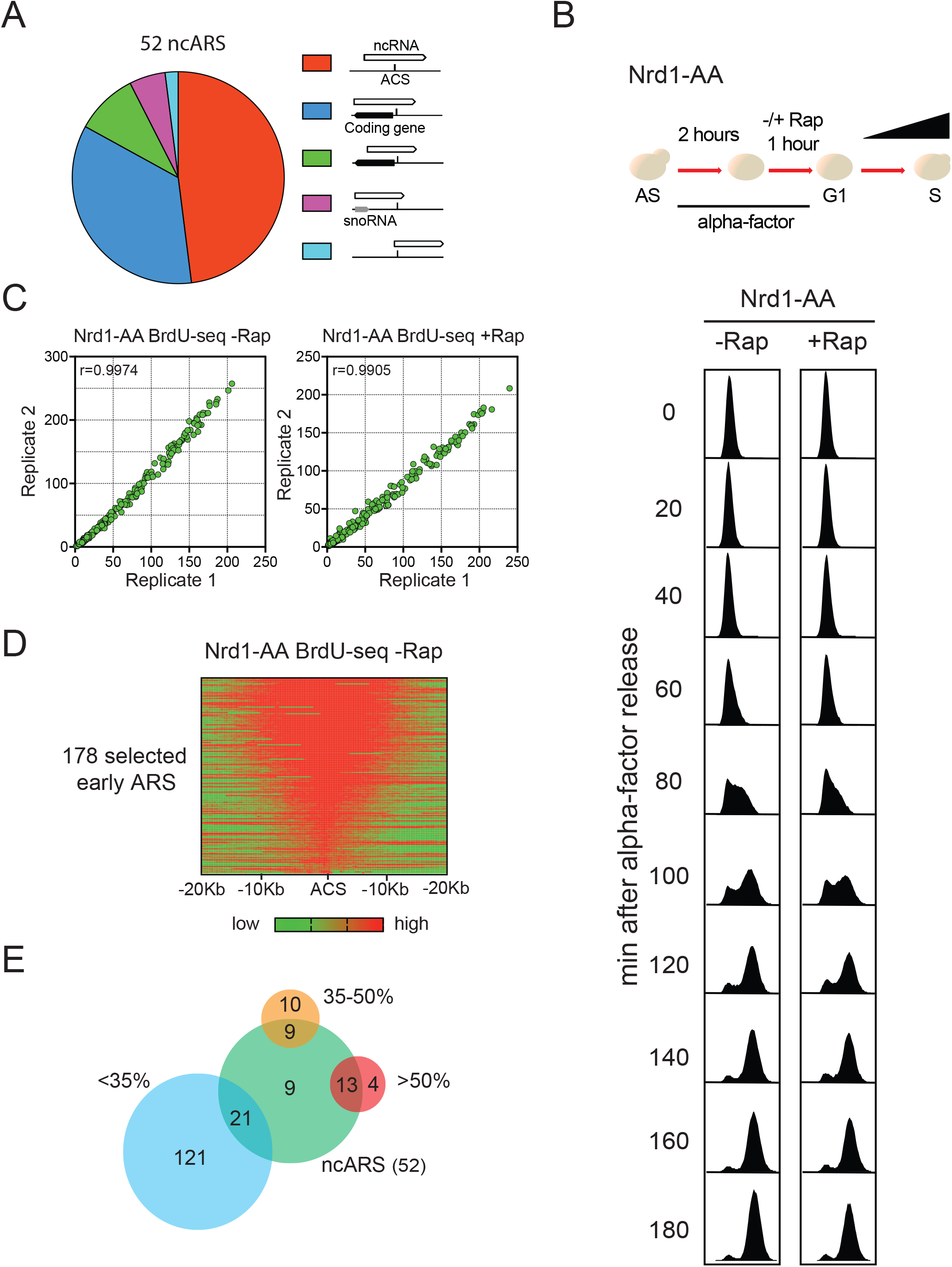
related to Figure 1. **(A)** Configuration distributions of the 52 ncRNAs-containing ARS (ncARS) with respect to the ACS. **(B)** FACS profiles of S-phase progression for the Nrd1-AA strain. **(C)** The mean coverage of BrdU nascent DNA in a 5Kb window centered around the ACS was measured for the 178 ARS taken into account in Figure 1E. The plots represent this mean coverage for the 2 replicates in either –Rap or +Rap. **(D)** Heatmap representing the BrdU-seq densities (log2) 20Kb around the ACS of the 178 replication origins considered as active in the experiment. **(E)** Overlap between the 52 ncRNA-containing ARS and the different classes of BrdU incorporation-defective replication origins in +Rap versus –Rap. The ncARS strongly overlap with the most affected ARS.

**Supplementary Figure 2:**
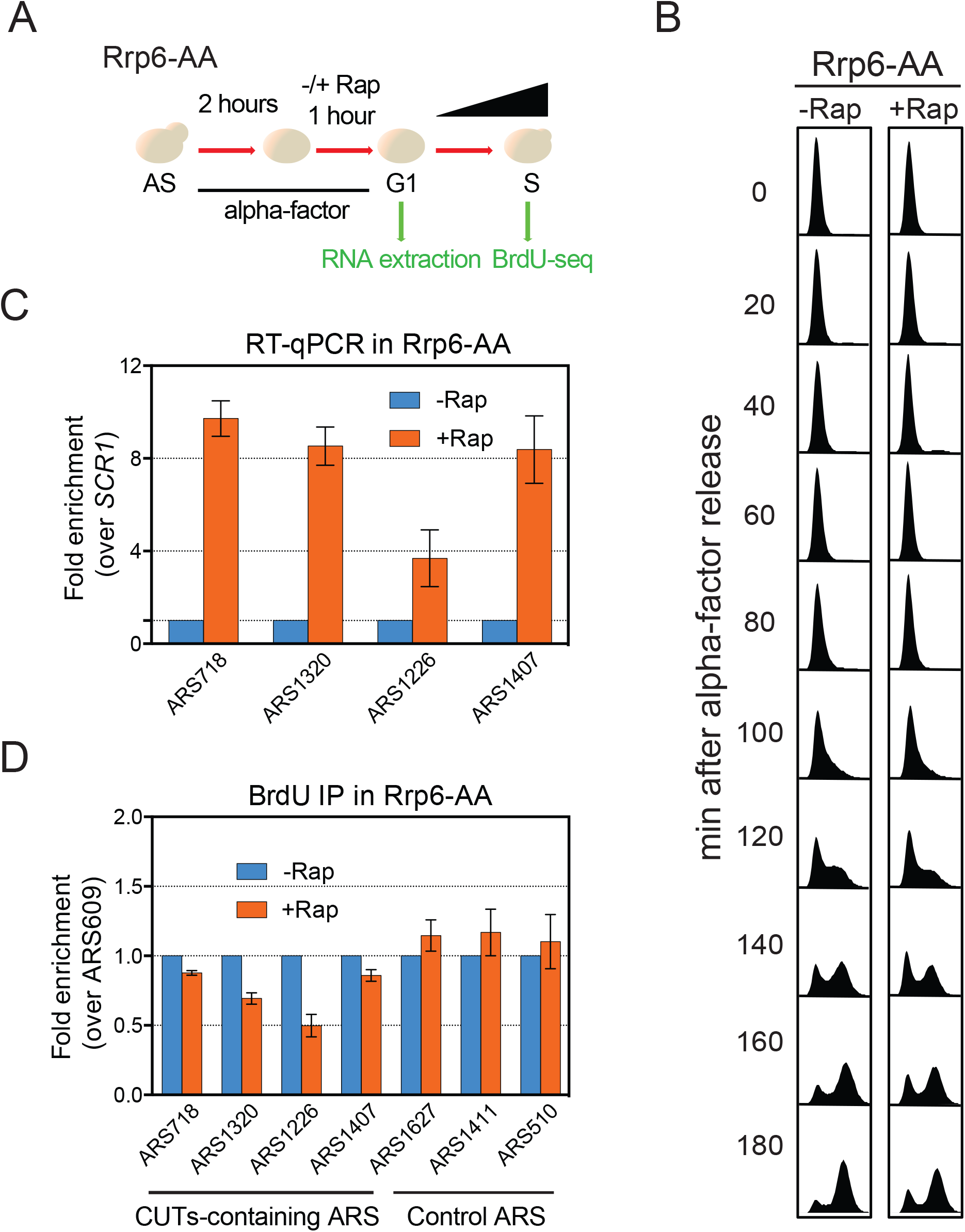
related to Figure 1. **(A)** Rrp6-AA cells were treated as in Figure 1C. RNA was extracted in G1-phase while DNA extraction and BrdU immunoprecipitation were performed at the 80 min time point following release into S-phase. **(B)** FACS profiles of S-phase progression for the Rrp6-AA strain. **(C)** RT-qPCR analysis of CUTs-containing ARS in G1-phase synchronized cells with or without 1h rapamycin treatment. Measured ncRNAs were normalized to *SCR1* RNA. **(D)** Analysis of BrdU incorporation after 80 min of S-phase release at 18°C. Three ARS which do not contain CUTs were used as controls. Fold enrichment represents the ratio of immunoprecipitated BrdU for a given ARS over the value of the very late replicated origin ARS609. For **(C)** and **(D)**, the fold enrichment was artificially set to 1 for the –Rap condition (n=3). Error bars represent SEM.

**Supplementary Figure 3:**
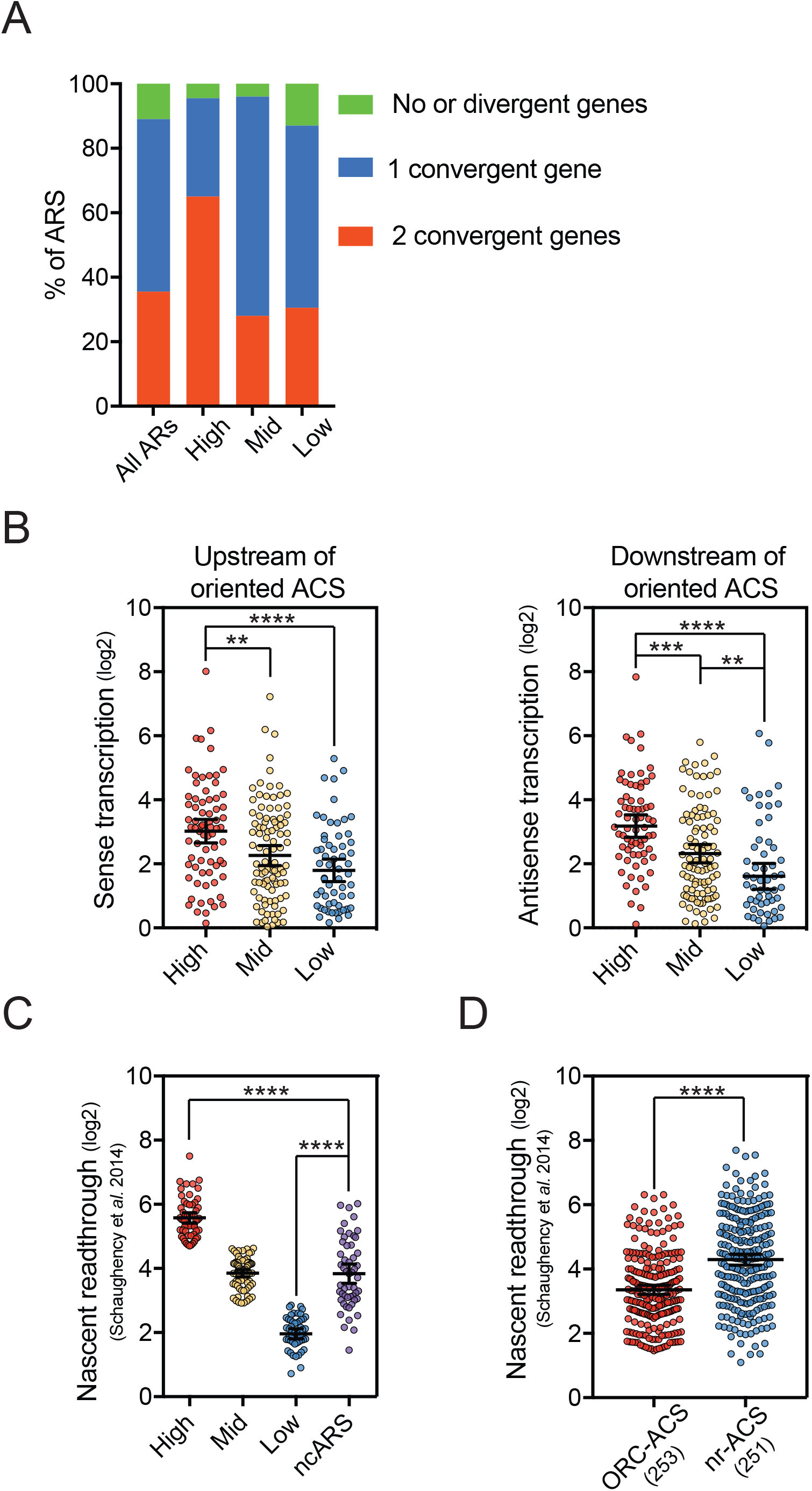
related to Figure 3. **(A)** Gene configuration around All ARS, High, Mid and Low classes of transcribed ARS. A gene was considered as convergent if pointing to the ARS within a distance <500bp to the ACS. **(B)** Scatter dot plot of nascent transcription pointing toward the ARS either Upstream (−400 to −300bp) or Downstream (+400 to +500bp) relative to the oriented ACS. **(C)** Scatter dot plot representing natural readthrough of CUTs/NUTs-containing ARS (ncARS) versus High, Mid and Low classes. **(D)** Scatter dot plot presenting nascent transcription in the ACS to +100bp region for ORC-bound ACS (ORC-ACS) and non-replicated ACS (nrACS) as defined in Eaton et al., 2010.

**Supplementary Figure 4:**
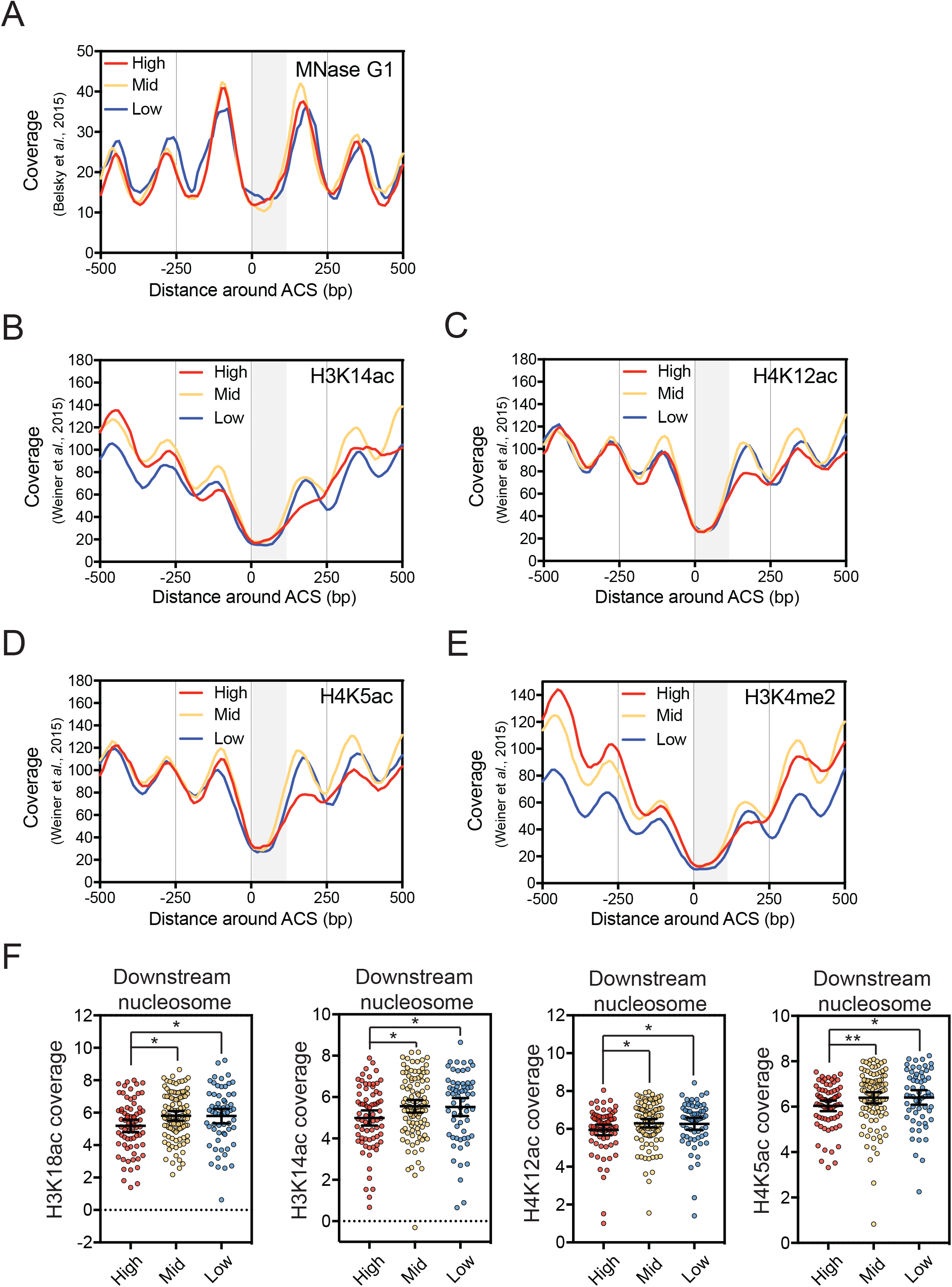
related to Figure 3. **(A)(B)(C)(D)(E)** Metagene analysis of G1-phase MNase-seq, H3K14ac, H4K12ac, H4K5ac and H3K4me2 profiles around the ACS of the High, Mid and Low classes of ARS. Data were retrieved from^16,47,48^. Results were smoothed over a 10bp-moving window. No significant differences were detected over the ACS to +100bp region. **(F)** H3K18ac, H3K14ac, H4K12ac, and H4K5ac coverages were measured over the downstream nucleosome (between 160bp to 200bp from the ACS) of the 3 classes considering this region as 1 bin.

**Supplementary Figure 5:**
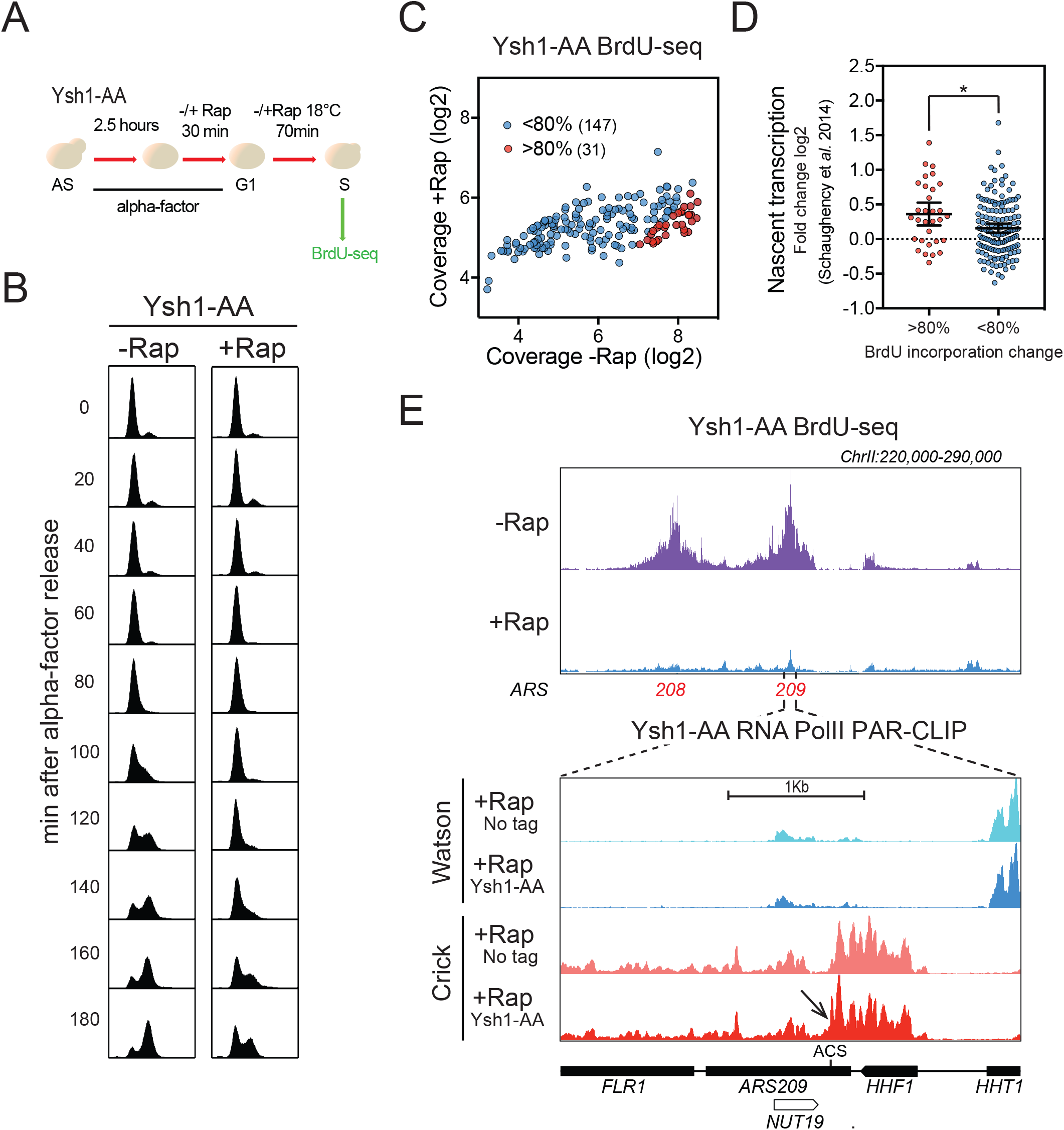
related to Figure 4. **(A)** Ysh1-AA cells were synchronized in G1-phase with alpha-factor during 3 hours at 30°C. During the last 30min, rapamycin (Rap) was added or not in the medium. Cells were then washed and released into the cell cycle at 18°C in the presence of BrdU and −/+Rap. After 70min, cells were collected for DNA extraction and BrdU-seq. **(B)** FACS analysis of the Ysh1-AA strain. Cells were treated as indicated in A. **(C)** Scatter plot depicting the mean coverage of BrdU nascent DNA over the same 178 ARS as in Figure 1E. The 31 red dots and 147 blue dots represent the ARS showing >80% and <80% decrease in BrdU incorporation in +Rap versus –Rap respectively. **(D)** Scatter dot-plots representing the ratio log2 +Rap/-Rap of the RNA PolII PAR-CLIP signal from both strands between the ACS and +100 of oriented ARS in the Ysh1-AA strain at the 2 classes of ARS defined above. **(E)** Top: Snapshot illustrating the BrdU incorporation defect in the Ysh1-AA strain. Bottom: zoom over the ARS209 showing the *HHF1* gene readthrough transcription over the ACS by RNA PolII PAR-CLIP. Transcriptional readthrough is indicated by an arrow.

**Supplementary Figure 6:**
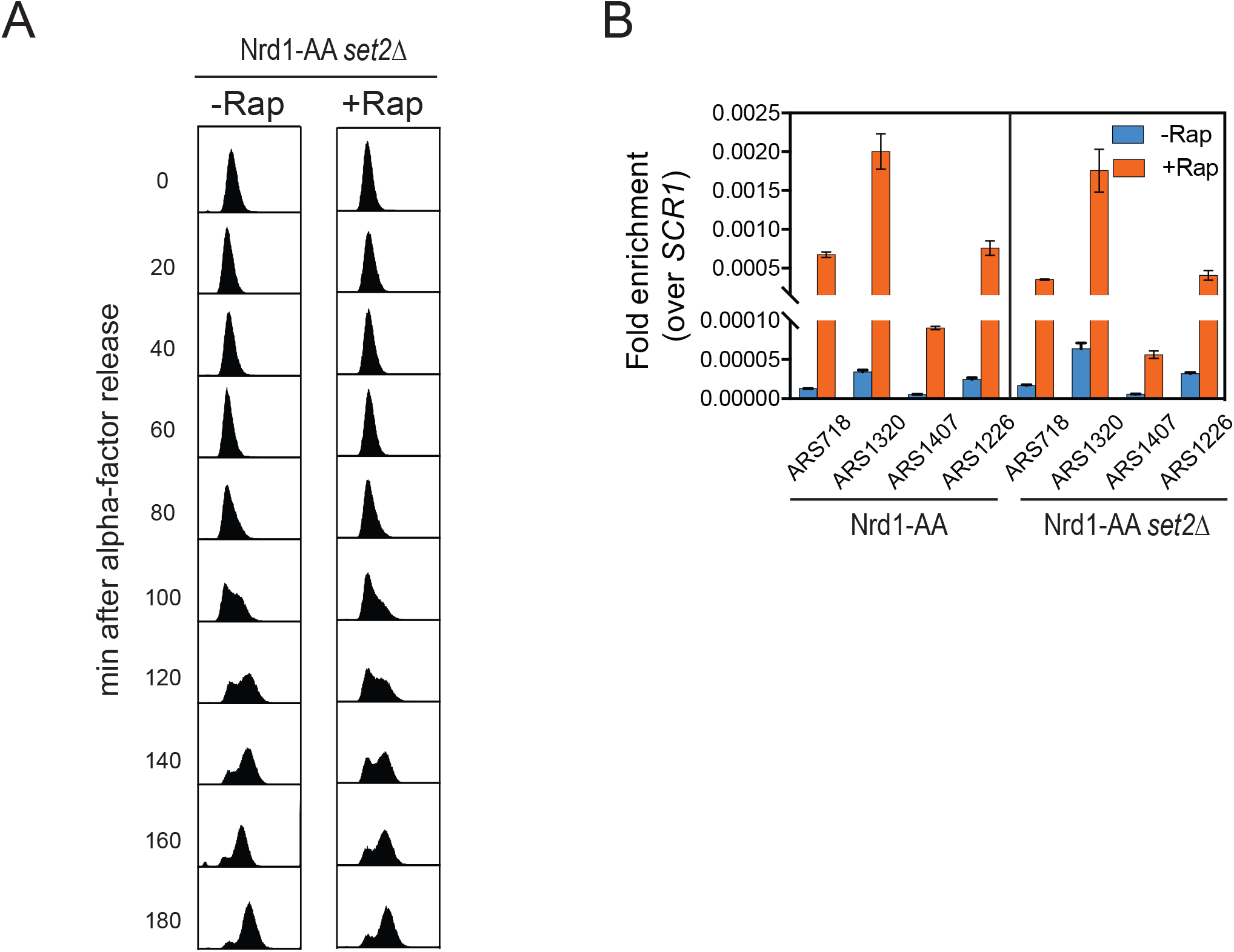
related to Figure 5. **(A)** FACS analysis of the Nrd1-AA *set2Δ* strain. Cells were treated as shown in Figure 5A. **(B)** RT-qPCR analysis of NUTs-containing ARS in the Nrd1-AA and Nrd1-AA *set2Δ* strains with or without 1h rapamycin treatment. Measured ncRNAs were normalized to *SCR1* RNA.

**Supplementary Table 1.** List of ARS features used in this study. The 178 ARS analyzed for BrdU incorporation (Figures 1, 4 and 5) are indicated in grey.

| **Chr** | **ACS** | **ARS** | **Orientation** | **Timing**<br>**(min)** | **Efficiency**<br>**(%)** |
| --- | --- | --- | --- | --- | --- |
| chrI | 147536 | ARS108 | - | 23.735 | 0.668 |
| chrI | 124521 | ARS107 | - | 27.404 | 0.741 |
| chrI | 70432 | ARS106 | - | 31.291 | 0.218 |
| chrI | 42059 | ARS105 | + | 28.660 | 0.326 |
| chrII | 237875 | ARS208 | - | 15.588 | 0.695 |
| chrII | 63373 | ARS202 | - | 19.066 | 0.616 |
| chrII | 486897 | ARS216 | - | 19.982 | 0.760 |
| chrII | 255081 | ARS209 | - | 24.969 | 0.501 |
| chrII | 418010 | ARS215 | - | 25.267 | 0.488 |
| chrII | 774020 | ARS227 | - | 27.550 | 0.431 |
| chrII | 802270 | ARS229 | - | 28.119 | 0.588 |
| chrII | 170263 | ARS207 | - | 31.714 | 0.377 |
| chrII | 408041 | ARS214 | - | 34.844 | 0.391 |
| chrII | 622754 | ARS220 | + | 21.272 | 0.700 |
| chrII | 326193 | ARS211 | + | 27.165 | 0.766 |
| chrII | 632018 | ARS221 | + | 29.128 | 0.301 |
| chrII | 611771 | ARS219.5 | + | 32.332 | 0.255 |
| chrIII | 39591 | ARS305 | - | 16.238 | 0.591 |
| chrIII | 108972 | ARS307 | - | 17.568 | 0.345 |
| chrIII | 74523 | ARS306 | + | 14.501 | 0.777 |
| chrIII | 224858 | ARS315 | + | 17.452 | 0.837 |
| chrIII | 166657 | ARS310 | + | 18.158 | 0.409 |
| chrIII | 132042 | ARS309 | + | 21.817 | 0.543 |
| chrIII | 315816 | ARS319 | + | 23.144 | 0.247 |
| chrIV | 484036 | ARS417 | - | 16.834 | 0.536 |
| chrIV | 555399 | ARS418 | - | 17.234 | 0.924 |
| chrIV | 1159443 | ARS432 | - | 18.662 | 0.478 |
| chrIV | 1166175 | ARS432.5 | - | 19.201 | 0.644 |
| chrIV | 1302759 | ARS435 | - | 20.016 | 0.704 |
| chrIV | 505519 | ARS417.5 | - | 20.838 | 0.562 |
| chrIV | 329742 | ARS413 | - | 21.937 | 0.660 |
| chrIV | 212593 | ARS409 | - | 29.550 | 0.558 |
| chrIV | 46222 | ARS404 | - | 30.060 | 0.668 |
| chrIV | 629309 | ARS420 | - | 31.775 | 0.587 |
| chrIV | 640065 | ARS421 | - | 34.626 | 0.415 |
| chrIV | 1461904 | ARS442 | + | 19.074 | 0.278 |
| chrIV | 408132 | ARS414 | + | 20.703 | 0.374 |
| chrIV | 913862 | ARS428 | + | 21.341 | 0.401 |
| chrIV | 921742 | ARS429 | + | 21.572 | 0.243 |
| chrIV | 316878 | ARS412 | + | 24.365 | 0.289 |
| chrIV | 123677 | ARS406 | + | 27.245 | 0.512 |
| chrIV | 1057893 | ARS431 | + | 29.587 | 0.506 |
| chrIX | 73953 | ARS907 | - | 19.888 | 0.350 |
| chrIX | 105966 | ARS909 | - | 21.972 | 0.790 |
| chrIX | 412000 | ARS922 | - | 22.149 | 0.660 |
| chrIX | 342028 | ARS919 | - | 27.051 | 0.579 |
| chrIX | 310739 | ARS916 | - | 30.660 | 0.557 |
| chrIX | 214733 | ARS913 | + | 15.871 | 0.643 |
| chrIX | 175171 | ARS912 | + | 26.345 | 0.628 |
| chrIX | 357223 | ARS920 | + | 26.473 | 0.604 |
| chrIX | 80373 | ARS908 | + | 28.215 | 0.413 |
| chrV | 173807 | ARS511 | - | 18.779 | 0.890 |
| chrV | 59469 | ARS507 | - | 19.620 | 0.798 |
| chrV | 145713 | ARS510 | - | 19.651 | 0.583 |
| chrV | 406902 | ARS517 | - | 19.977 | 0.881 |
| chrV | 94056 | ARS508 | + | 17.251 | 0.914 |
| chrV | 353583 | ARS516 | + | 20.327 | 0.833 |
| chrV | 569630 | ARS523 | + | 21.305 | 0.403 |
| chrV | 287566 | ARS514 | + | 26.321 | 0.502 |
| chrV | 212455 | ARS512 | + | 26.816 | 0.727 |
| chrV | 498900 | ARS520 | + | 27.741 | 0.758 |
| chrV | 549586 | ARS522 | + | 30.072 | 0.372 |
| chrVI | 167731 | ARS606 | - | 20.287 | 0.756 |
| chrVI | 118678 | ARS603.5 | - | 26.961 | 0.556 |
| chrVI | 136037 | ARS605 | - | 36.277 | 0.247 |
| chrVI | 199402 | ARS607 | + | 15.378 | 0.936 |
| chrVI | 127869 | ARS604 | + | 19.365 | 0.299 |
| chrVI | 68832 | ARS603 | + | 25.803 | 0.659 |
| chrVII | 485115 | ARS719 | - | 16.890 | 0.771 |
| chrVII | 834669 | ARS731 | - | 17.174 | 0.860 |
| chrVII | 508911 | ARS720 | - | 19.346 | 0.464 |
| chrVII | 653836 | ARS726 | - | 19.528 | 0.603 |
| chrVII | 388849 | ARS717 | - | 20.561 | 0.773 |
| chrVII | 977910 | ARS733 | - | 21.676 | 0.608 |
| chrVII | 421285 | ARS718 | - | 21.747 | 0.825 |
| chrVII | 352866 | ARS716 | - | 25.985 | 0.717 |
| chrVII | 64458 | ARS702 | - | 30.001 | 0.633 |
| chrVII | 660004 | ARS727 | - | 31.979 | 0.335 |
| chrVII | 17906 | ARS700.5 | - | 33.251 | 0.294 |
| chrVII | 1083677 | ARS131a | + | 17.176 | 0.268 |
| chrVII | 888418 | ARS731.5 | + | 17.603 | 0.786 |
| chrVII | 715319 | ARS728 | + | 21.386 | 0.803 |
| chrVII | 163242 | ARS707 | + | 23.729 | 0.666 |
| chrVII | 203978 | ARS710 | + | 32.839 | 0.466 |
| chrVIII | 392250 | ARS818 | - | 20.133 | 0.462 |
| chrVIII | 501949 | ARS822 | - | 20.436 | 0.657 |
| chrVIII | 447794 | ARS820 | - | 20.891 | 0.891 |
| chrVIII | 359698 | ARS816 | - | 21.387 | 0.699 |
| chrVIII | 45778 | ARS804 | - | 27.409 | 0.216 |
| chrVIII | 297102 | ARS815 | + | 19.650 | 0.908 |
| chrVIII | 64301 | ARS805 | + | 21.111 | 0.637 |
| chrVIII | 168599 | ARS809 | + | 23.838 | 0.563 |
| chrVIII | 245791 | ARS813 | + | 25.611 | 0.834 |
| chrVIII | 556152 | ARS824 | + | 38.395 | 0.163 |
| chrX | 417311 | ARS1014 | - | 18.061 | 0.769 |
| chrX | 654465 | ARS1020 | - | 20.441 | 0.483 |
| chrX | 442416 | ARS1015 | - | 22.269 | 0.271 |
| chrX | 113736 | ARS1007 | - | 22.705 | 0.662 |
| chrX | 67713 | ARS1005 | - | 24.249 | 0.834 |
| chrX | 228567 | ARS1009 | - | 26.895 | 0.449 |
| chrX | 374861 | ARS1012 | + | 16.843 | 0.790 |
| chrX | 683928 | ARS1021 | + | 21.242 | 0.836 |
| chrX | 161448 | ARS1007.5 | + | 22.882 | 0.679 |
| chrXI | 55867 | ARS1103 | - | 20.460 | 0.859 |
| chrXI | 329502 | ARS1109 | - | 23.609 | 0.669 |
| chrXI | 213310 | ARS1106.7 | - | 24.563 | 0.611 |
| chrXI | 153125 | ARS1106 | - | 24.774 | 0.742 |
| chrXI | 457168 | ARS1114.5 | - | 26.925 | 0.386 |
| chrXI | 612053 | ARS1120 | - | 27.419 | 0.660 |
| chrXI | 388668 | ARS1112 | + | 23.775 | 0.708 |
| chrXI | 98390 | ARS1104.5 | + | 23.943 | 0.806 |
| chrXI | 257590 | ARS1107 | + | 24.564 | 0.823 |
| chrXI | 642422 | ARS1123 | + | 24.832 | 0.285 |
| chrXI | 416884 | ARS1113 | + | 27.822 | 0.508 |
| chrXII | 373328 | ARS1213 | - | 16.165 | 0.924 |
| chrXII | 513085 | ARS1217 | - | 18.729 | 0.929 |
| chrXII | 412853 | ARS1215 | - | 20.842 | 0.791 |
| chrXII | 794192 | ARS1226 | - | 21.585 | 0.724 |
| chrXII | 231251 | ARS1211 | + | 18.443 | 0.886 |
| chrXII | 1007238 | ARS1232 | + | 22.050 | 0.653 |
| chrXII | 659895 | ARS1220 | + | 23.299 | 0.616 |
| chrXII | 730540 | ARS1222 | + | 24.034 | 0.617 |
| chrXII | 243743 | ARS1211.5 | + | 31.538 | 0.195 |
| chrXII | 622861 | ARS1219 | + | 31.913 | 0.218 |
| chrXII | 1013787 | ARS1233 | + | 34.939 | 0.206 |
| chrXIII | 503627 | ARS1319 | - | 18.955 | 0.693 |
| chrXIII | 535769 | ARS1320 | - | 20.864 | 0.788 |
| chrXIII | 94390 | ARS1305 | - | 22.511 | 0.759 |
| chrXIII | 554598 | ARS1322 | - | 23.191 | 0.351 |
| chrXIII | 897975 | ARS1332 | - | 25.787 | 0.812 |
| chrXIII | 40132 | ARS1304 | - | 30.096 | 0.243 |
| chrXIII | 758416 | ARS1327 | - | 31.011 | 0.569 |
| chrXIII | 815391 | ARS1330 | + | 18.728 | 0.702 |
| chrXIII | 31767 | ARS1303 | + | 20.124 | 0.346 |
| chrXIII | 263127 | ARS1309 | + | 20.780 | 0.589 |
| chrXIII | 634521 | ARS1324 | + | 21.295 | 0.449 |
| chrXIII | 433030 | ARS1315 | + | 21.650 | 0.590 |
| chrXIII | 137322 | ARS1307 | + | 21.664 | 0.675 |
| chrXIII | 805162 | ARS1329 | + | 23.111 | 0.410 |
| chrXIII | 371020 | ARS1312 | + | 23.242 | 0.832 |
| chrXIII | 649362 | ARS1325 | + | 23.833 | 0.570 |
| chrXIII | 159062 | ARS1307.5 | + | 27.358 | 0.438 |
| chrXIII | 611318 | ARS1323 | + | 33.987 | 0.479 |
| chrXIV | 89756 | ARS1407 | - | 22.498 | 0.673 |
| chrXIV | 280067 | ARS1414 | - | 23.015 | 0.690 |
| chrXIV | 691682 | ARS1427 | - | 23.675 | 0.672 |
| chrXIV | 635835 | ARS1426 | - | 26.400 | 0.509 |
| chrXIV | 412443 | ARS1417 | - | 27.879 | 0.727 |
| chrXIV | 546150 | ARS1421 | - | 28.955 | 0.352 |
| chrXIV | 449538 | ARS1419 | - | 30.269 | 0.616 |
| chrXIV | 322006 | ARS1415 | + | 20.947 | 0.780 |
| chrXIV | 609538 | ARS1424 | + | 23.739 | 0.779 |
| chrXIV | 499042 | ARS1420 | + | 25.434 | 0.725 |
| chrXIV | 61696 | ARS1406 | + | 28.945 | 0.584 |
| chrXV | 277733 | ARS1511 | - | 19.353 | 0.889 |
| chrXV | 309249 | ARS1512 | - | 19.762 | 0.424 |
| chrXV | 874369 | ARS1526 | - | 22.955 | 0.616 |
| chrXV | 155258 | ARS1509.5 | - | 28.308 | 0.165 |
| chrXV | 337484 | ARS1513 | - | 31.274 | 0.389 |
| chrXV | 85360 | ARS1508 | - | 31.364 | 0.341 |
| chrXV | 167004 | ARS1510 | + | 21.679 | 0.788 |
| chrXV | 113910 | ARS1509 | + | 22.066 | 0.746 |
| chrXV | 766691 | ARS1523 | + | 23.198 | 0.767 |
| chrXV | 436793 | ARS1513.5 | + | 24.732 | 0.610 |
| chrXV | 72689 | ARS1507 | + | 25.645 | 0.581 |
| chrXV | 908311 | ARS1528 | + | 33.991 | 0.278 |
| chrXV | 35714 | ARS1506.5 | + | 34.024 | 0.199 |
| chrXVI | 777096 | ARS1626.5 | - | 17.945 | 0.838 |
| chrXVI | 43150 | ARS1604 | - | 19.931 | 0.170 |
| chrXVI | 695620 | ARS1625 | - | 25.280 | 0.689 |
| chrXVI | 842852 | ARS1628 | - | 33.042 | 0.415 |
| chrXVI | 289531 | ARS1614 | + | 18.272 | 0.623 |
| chrXVI | 73105 | ARS1605 | + | 20.350 | 0.745 |
| chrXVI | 384595 | ARS1618 | + | 20.991 | 0.560 |
| chrXVI | 633924 | ARS1623 | + | 21.153 | 0.895 |
| chrXVI | 511707 | ARS1621 | + | 22.719 | 0.693 |
| chrXVI | 116594 | ARS1607 | + | 23.358 | 0.636 |
| chrXVI | 684408 | ARS1624 | + | 31.888 | 0.333 |
| chrI | 215011 | ARS111 | - | 18.780 | 0.670 |
| chrI | 6572 | ARS201 | - | 26.714 | 0.379 |
| chrII | 539400 | ARS218 | - | 18.339 | 0.356 |
| chrII | 517285 | ARS217 | - | 22.380 | 0.192 |
| chrII | 143692 | ARS206 | - | 26.351 | 0.586 |
| chrII | 741781 | ARS224 | - | 28.218 | 0.521 |
| chrII | 591434 | ARS219 | + | 18.575 | 0.447 |
| chrII | 93553 | ARS203 | + | 22.068 | 0.315 |
| chrII | 379119 | ARS212 | + | 26.964 | 0.386 |
| chrII | 757485 | ARS225 | + | 30.997 | 0.319 |
| chrIII | 273027 | ARS316 | - | 22.416 | 0.465 |
| chrIV | 1033263 | ARS430.5 | - | 16.136 | 0.434 |
| chrIV | 236050 | ARS409.5 | - | 20.600 | 0.681 |
| chrIV | 101241 | ARS405.5 | - | 21.706 | 0.471 |
| chrIV | 1487095 | ARS446 | - | 24.132 | 0.454 |
| chrIV | 852 | ARS400 | - | 25.988 | 0.566 |
| chrIV | 1110136 | ARS431.5 | - | 27.711 | 0.629 |
| chrIV | 86124 | ARS405 | - | 42.903 | 0.189 |
| chrIV | 702929 | ARS422 | + | 21.481 | 0.701 |
| chrIV | 1404327 | ARS440 | + | 22.368 | 0.489 |
| chrIV | 1276272 | ARS434 | + | 27.188 | 0.651 |
| chrIV | 1240924 | ARS433 | + | 27.513 | 0.820 |
| chrIV | 845128 | ARS426.5 | + | 28.403 | 0.359 |
| chrIV | 253841 | ARS410 | + | 29.972 | 0.183 |
| chrIX | 136287 | ARS911 | - | 22.478 | 0.561 |
| chrV | 520952 | ARS521 | + | 30.851 | 0.392 |
| chrVII | 187407 | ARS709 | + | 28.594 | 0.394 |
| chrVII | 1063520 | ARS735.5 | + | 33.917 | 0.580 |
| chrVIII | 535621 | ARS823.5 | - | 28.884 | 0.540 |
| chrX | 730035 | ARS1023 | + | 17.486 | 0.196 |
| chrX | 337270 | ARS1011 | + | 20.057 | 0.494 |
| chrX | 298838 | ARS1010 | + | 25.888 | 0.678 |
| chrX | 23928 | ARS1004 | + | 48.747 | 0.337 |
| chrXII | 52108 | ARS1203 | - | 24.886 | 0.337 |
| chrXII | 888740 | ARS1227.5 | - | 25.101 | 0.768 |
| chrXII | 289421 | ARS1212 | - | 28.695 | 0.684 |
| chrXII | 1024153 | ARS1234 | - | 29.394 | 0.409 |
| chrXII | 822105 | ARS1227 | + | 28.461 | 0.668 |
| chrXIII | 468237 | ARS1316 | + | 23.344 | 0.441 |
| chrXIII | 688918 | ARS1326 | + | 28.514 | 0.392 |
| chrXIII | 878735 | ARS1331.7 | + | 30.640 | 0.262 |
| chrXIII | 772677 | ARS1328 | + | 32.178 | 0.310 |
| chrXIV | 250465 | ARS1413 | - | 25.503 | 0.530 |
| chrXIV | 764337 | ARS1429 | - | 26.026 | 0.416 |
| chrXIV | 169749 | ARS1411 | - | 27.602 | 0.622 |
| chrXIV | 196226 | ARS1412 | - | 31.148 | 0.626 |
| chrXIV | 126679 | ARS1410 | - | 37.763 | 0.342 |
| chrXIV | 738730 | ARS1428.5 | + | 35.238 | 0.442 |
| chrXV | 566598 | ARS1516 | - | 21.689 | 0.780 |
| chrXV | 681350 | ARS1520 | - | 29.163 | 0.622 |
| chrXV | 729797 | ARS1521 | + | 19.448 | 0.316 |
| chrXV | 783388 | ARS1524 | + | 22.563 | 0.346 |
| chrXV | 656703 | ARS1519 | + | 26.471 | 0.632 |
| chrXVI | 584397 | ARS1622.7 | - | 29.081 | 0.259 |
| chrXVI | 331771 | ARS1617 | + | 23.235 | 0.477 |
| chrXVI | 880921 | ARS1630 | + | 30.530 | 0.613 |

**Supplementary Table 2.** List of strains used in this study

| **Strain** | **Lab name** | **Genotype** |
| --- | --- | --- |
| Nrd1-AA | FSY5348 | *MAT a, tor1-1, fpr1::NAT, RPL13A-2×FKBP12::TRP1, NRD1-FRB::KanMX6, bar1::LEU2, BrdU-inc::HIS3, ura3-1, ade2-1* |
| Nrd1-AA *set2Δ* | FSY7284 | *MAT a, tor1-1, fpr1::NAT, RPL13A-2×FKBP12::TRP1, NRD1-FRB::KanMX6, bar1::LEU2, BrdU-inc::HIS3, set2 ::URA3, ade2-1* |
| Rrp6-AA | FSY5725 | *MAT a, tor1-1, fpr1::loxP-LEU2-loxP, RPL13A-2×FKBP12::loxP, RRP6-FRB::KanMX6, bar1::URA3, BrdU-inc::HIS3, ade 2-1 trp1-1, leu2-3,112* |
| Ysh1-AA | FSY7715 | *MAT a, tor1-1, fpr1::loxP-LEU2-loxP, RPL13A-2×FKBP12::loxP, YSH1-FRB::KanMX6, bar1::URA3, BrdU-inc::HIS3, ade 2-1 trp1-1, leu2-3,112* |

## REFERENCES

1. Nieduszynski, C.A., Knox, Y. & Donaldson, A.D. Genome-wide identification of replication origins in yeast by comparative genomics. Genes Dev 20, 1874–9 (2006).

2. Stinchcomb, D.T., Struhl, K. & Davis, R.W. Isolation and characterisation of a yeast chromosomal replicator. Nature 282, 39–43 (1979).

3. Bell, S.P. Eukaryotic replicators and associated protein complexes. Curr Opin Genet Dev 5, 162–7 (1995).

4. Eaton, M.L., Galani, K., Kang, S., Bell, S.P. & MacAlpine, D.M. Conserved nucleosome positioning defines replication origins. Genes Dev 24, 748–53 (2010).

5. Segal, E. & Widom, J. Poly(dA:dT) tracts: major determinants of nucleosome organization. Curr Opin Struct Biol 19, 65–71 (2009).

6. Hawkins, M. et al. High-resolution replication profiles define the stochastic nature of genome replication initiation and termination. Cell Rep 5, 1132–41 (2013).

7. McGuffee, S.R., Smith, D.J. & Whitehouse, I. Quantitative, genome-wide analysis of eukaryotic replication initiation and termination. Molecular cell 50, 123–35 (2013).

8. Raghuraman, M.K. et al. Replication dynamics of the yeast genome. Science 294, 115–21 (2001).

9. Deegan, T.D. & Diffley, J.F. MCM: one ring to rule them all. Curr Opin Struct Biol 37, 145–51 (2016).

10. Aparicio, O.M. Location, location, location: it’s all in the timing for replication origins. Genes Dev 27, 117–28 (2013).

11. Czajkowsky, D.M., Liu, J., Hamlin, J.L. & Shao, Z. DNA combing reveals intrinsic temporal disorder in the replication of yeast chromosome VI. J Mol Biol 375, 12–9 (2008).

12. Yang, S.C., Rhind, N. & Bechhoefer, J. Modeling genome-wide replication kinetics reveals a mechanism for regulation of replication timing. Mol Syst Biol 6, 404 (2010).

13. Yoshida, K., Poveda, A. & Pasero, P. Time to be versatile: regulation of the replication timing program in budding yeast. J Mol Biol 425, 4696–705 (2013).

14. Soriano, I., Morafraile, E.C., Vazquez, E., Antequera, F. & Segurado, M. Different nucleosomal architectures at early and late replicating origins in Saccharomyces cerevisiae. BMC Genomics 15, 791 (2014).

15. Rodriguez, J., Lee, L., Lynch, B. & Tsukiyama, T. Nucleosome occupancy as a novel chromatin parameter for replication origin functions. Genome Res 27, 269–277 (2017).

16. Belsky, J.A., MacAlpine, H.K., Lubelsky, Y., Hartemink, A.J. & MacAlpine, D.M. Genome-wide chromatin footprinting reveals changes in replication origin architecture induced by pre-RC assembly. Genes & development 29, 212–24 (2015).

17. Unnikrishnan, A., Gafken, P.R. & Tsukiyama, T. Dynamic changes in histone acetylation regulate origins of DNA replication. Nature structural & molecular biology 17, 430–7 (2010).

18. Vogelauer, M., Rubbi, L., Lucas, I., Brewer, B.J. & Grunstein, M. Histone acetylation regulates the time of replication origin firing. Molecular cell 10, 1223–33 (2002).

19. Knott, S.R., Viggiani, C.J., Tavare, S. & Aparicio, O.M. Genome-wide replication profiles indicate an expansive role for Rpd3L in regulating replication initiation timing or efficiency, and reveal genomic loci of Rpd3 function in Saccharomyces cerevisiae. Genes Dev 23, 1077–90 (2009).

20. Aparicio, J.G., Viggiani, C.J., Gibson, D.G. & Aparicio, O.M. The Rpd3-Sin3 histone deacetylase regulates replication timing and enables intra-S origin control in Saccharomyces cerevisiae. Mol Cell Biol 24, 4769–80 (2004).

21. Fraser, H.B. Cell-cycle regulated transcription associates with DNA replication timing in yeast and human. Genome Biol 14, R111 (2013).

22. Knott, S.R. et al. Forkhead transcription factors establish origin timing and long-range clustering in S. cerevisiae. Cell 148, 99–111 (2012).

23. Mayan, M.D. RNAP-II molecules participate in the anchoring of the ORC to rDNA replication origins. PLoS One 8, e53405 (2013).

24. Mori, S. & Shirahige, K. Perturbation of the activity of replication origin by meiosis-specific transcription. J Biol Chem 282, 4447–52 (2007).

25. Blitzblau, H.G., Chan, C.S., Hochwagen, A. & Bell, S.P. Separation of DNA replication from the assembly of break-competent meiotic chromosomes. PLoS genetics 8, e1002643 (2012).

26. Snyder, M., Sapolsky, R.J. & Davis, R.W. Transcription interferes with elements important for chromosome maintenance in Saccharomyces cerevisiae. Molecular and cellular biology 8, 2184–94 (1988).

27. Gros, J. et al. Post-licensing Specification of Eukaryotic Replication Origins by Facilitated Mcm2-7 Sliding along DNA. Mol Cell 60, 797–807 (2015).

28. Nieduszynski, C.A., Blow, J.J. & Donaldson, A.D. The requirement of yeast replication origins for pre-replication complex proteins is modulated by transcription. Nucleic Acids Res 33, 2410–20 (2005).

29. Churchman, L.S. & Weissman, J.S. Nascent transcript sequencing visualizes transcription at nucleotide resolution. Nature 469, 368–73 (2011).

30. Schaughency, P., Merran, J. & Corden, J.L. Genome-wide mapping of yeast RNA polymerase II termination. PLoS Genet 10, e1004632 (2014).

31. David, L. et al. A high-resolution map of transcription in the yeast genome. Proc Natl Acad Sci U S A 103, 5320–5 (2006).

32. Arigo, J.T., Eyler, D.E., Carroll, K.L. & Corden, J.L. Termination of cryptic unstable transcripts is directed by yeast RNA-binding proteins Nrd1 and Nab3. Mol Cell 23, 841–51 (2006).

33. Steinmetz, E.J., Conrad, N.K., Brow, D.A. & Corden, J.L. RNA-binding protein Nrd1 directs poly(A)-independent 3’-end formation of RNA polymerase II transcripts. Nature 413, 327–31 (2001).

34. Tudek, A. et al. Molecular basis for coordinating transcription termination with noncoding RNA degradation. Mol Cell 55, 467–81 (2014).

35. Jensen, T.H., Jacquier, A. & Libri, D. Dealing with pervasive transcription. Mol Cell 52, 473–84 (2013).

36. Neil, H. et al. Widespread bidirectional promoters are the major source of cryptic transcripts in yeast. Nature 457, 1038–42 (2009).

37. van Dijk, E.L. et al. XUTs are a class of Xrn1-sensitive antisense regulatory non-coding RNA in yeast. Nature 475, 114–7 (2011).

38. Wyers, F. et al. Cryptic pol II transcripts are degraded by a nuclear quality control pathway involving a new poly(A) polymerase. Cell 121, 725–37 (2005).

39. Xu, Z. et al. Bidirectional promoters generate pervasive transcription in yeast. Nature 457, 1033–7 (2009).

40. Schulz, D. et al. Transcriptome surveillance by selective termination of noncoding RNA synthesis. Cell 155, 1075–87 (2013).

41. Porrua, O., Boudvillain, M. & Libri, D. Transcription Termination: Variations on Common Themes. Trends Genet 32, 508–22 (2016).

42. Baejen, C. et al. Genome-wide Analysis of RNA Polymerase II Termination at Protein-Coding Genes. Molecular cell (2017).

43. Kim, M. et al. The yeast Rat1 exonuclease promotes transcription termination by RNA polymerase II. Nature 432, 517–22 (2004).

44. Luo, W., Johnson, A.W. & Bentley, D.L. The role of Rat1 in coupling mRNA 3’-end processing to transcription termination: implications for a unified allosteric-torpedo model. Genes Dev 20, 954–65 (2006).

45. Haruki, H., Nishikawa, J. & Laemmli, U.K. The anchor-away technique: rapid, conditional establishment of yeast mutant phenotypes. Mol Cell 31, 925–32 (2008).

46. Castelnuovo, M. et al. Role of histone modifications and early termination in pervasive transcription and antisense-mediated gene silencing in yeast. Nucleic Acids Res 42, 4348–62 (2014).

47. Kubik, S. et al. Nucleosome Stability Distinguishes Two Different Promoter Types at All Protein-Coding Genes in Yeast. Mol Cell 60, 422–34 (2015).

48. Weiner, A. et al. High-resolution chromatin dynamics during a yeast stress response. Mol Cell 58, 371–86 (2015).

49. Rundlett, S.E. et al. HDA1 and RPD3 are members of distinct yeast histone deacetylase complexes that regulate silencing and transcription. Proc Natl Acad Sci U S A 93, 14503–8 (1996).

50. Woo, H., Dam Ha, S., Lee, S.B., Buratowski, S. & Kim, T. Modulation of gene expression dynamics by co-transcriptional histone methylations. Exp Mol Med 49, e326 (2017).

51. Mori, S. & Shirahige, K. Perturbation of the activity of replication origin by meiosis-specific transcription. The Journal of biological chemistry 282, 4447–52 (2007).

52. van Nues, R. et al. Kinetic CRAC uncovers a role for Nab3 in determining gene expression profiles during stress. Nat Commun 8, 12 (2017).

53. Bresson, S., Tuck, A., Staneva, D. & Tollervey, D. Nuclear RNA Decay Pathways Aid Rapid Remodeling of Gene Expression in Yeast. Mol Cell 65, 787–800 e5 (2017).

54. Wittmann, S. et al. The conserved protein Seb1 drives transcription termination by binding RNA polymerase II and nascent RNA. Nat Commun 8, 14861 (2017).

55. Heichinger, C., Penkett, C.J., Bahler, J. & Nurse, P. Genome-wide characterization of fission yeast DNA replication origins. EMBO J 25, 5171–9 (2006).

56. Hoggard, T., Shor, E., Muller, C.A., Nieduszynski, C.A. & Fox, C.A. A Link between ORC-origin binding mechanisms and origin activation time revealed in budding yeast. PLoS Genet 9, e1003798 (2013).

57. Das, S.P. et al. Replication timing is regulated by the number of MCMs loaded at origins. Genome Res 25, 1886–92 (2015).

58. Looke, M., Kristjuhan, K., Varv, S. & Kristjuhan, A. Chromatin-dependent and -independent regulation of DNA replication origin activation in budding yeast. EMBO Rep 14, 191–8 (2013).

59. Peace, J.M., Villwock, S.K., Zeytounian, J.L., Gan, Y. & Aparicio, O.M. Quantitative BrdU immunoprecipitation method demonstrates that Fkh1 and Fkh2 are rate-limiting activators of replication origins that reprogram replication timing in G1 phase. Genome Res 26, 365–75 (2016).

60. Pryde, F. et al. H3 k36 methylation helps determine the timing of cdc45 association with replication origins. PLoS One 4, e5882 (2009).

61. Carrozza, M.J. et al. Histone H3 methylation by Set2 directs deacetylation of coding regions by Rpd3S to suppress spurious intragenic transcription. Cell 123, 581–92 (2005).

62. Yoshida, K. et al. The histone deacetylases sir2 and rpd3 act on ribosomal DNA to control the replication program in budding yeast. Molecular cell 54, 691–7 (2014).

63. Malabat, C., Feuerbach, F., Ma, L., Saveanu, C. & Jacquier, A. Quality control of transcription start site selection by nonsense-mediated-mRNA decay. eLife 4 (2015).

64. Sequeira-Mendes, J. et al. Transcription initiation activity sets replication origin efficiency in mammalian cells. PLoS Genet 5, e1000446 (2009).

65. Dellino, G.I. et al. Genome-wide mapping of human DNA-replication origins: levels of transcription at ORC1 sites regulate origin selection and replication timing. Genome Res 23, 1–11 (2013).

66. Miotto, B., Ji, Z. & Struhl, K. Selectivity of ORC binding sites and the relation to replication timing, fragile sites, and deletions in cancers. Proc Natl Acad Sci U S A 113, E4810–9 (2016).

67. Preker, P. et al. RNA exosome depletion reveals transcription upstream of active human promoters. Science 322, 1851–4 (2008).

68. Nojima, T. et al. Mammalian NET-Seq Reveals Genome-wide Nascent Transcription Coupled to RNA Processing. Cell 161, 526–40 (2015).

69. Mayer, A. et al. Native elongating transcript sequencing reveals human transcriptional activity at nucleotide resolution. Cell 161, 541–54 (2015).

70. Petryk, N. et al. Replication landscape of the human genome. Nat Commun 7, 10208 (2016).

71. Viggiani, C.J. & Aparicio, O.M. New vectors for simplified construction of BrdU-Incorporating strains of Saccharomyces cerevisiae. Yeast 23, 1045–51 (2006).

72. Soudet, J., Jolivet, P. & Teixeira, M.T. Elucidation of the DNA end-replication problem in Saccharomyces cerevisiae. Molecular cell 53, 954–64 (2014).

73. David, F.P. et al. HTSstation: a web application and open-access libraries for high-throughput sequencing data analysis. PLoS One 9, e85879 (2014).

74. Camblong, J., Iglesias, N., Fickentscher, C., Dieppois, G. & Stutz, F. Antisense RNA stabilization induces transcriptional gene silencing via histone deacetylation in S. cerevisiae. Cell 131, 706–17 (2007).

75. Uwimana, N., Collin, P., Jeronimo, C., Haibe-Kains, B. & Robert, F. Bidirectional terminators in Saccharomyces cerevisiae prevent cryptic transcription from invading neighboring genes. Nucleic Acids Res 45, 6417–6426 (2017).

